# Development of a live-cell imaging assay to elucidate spatiotemporal dynamics of extracellular vesicle fusion with target cells

**DOI:** 10.1101/2025.07.23.666255

**Authors:** Jasper van den Ende, Kyra A.Y. Defourny, Huib H. Rabouw, Marvin E. Tanenbaum, Richard W. Wubbolts, Esther N.M. Nolte-‘t Hoen

## Abstract

Cells communicate via extracellular vesicles (EVs) containing functional RNAs, proteins and lipids. Knowledge on the fate of internalized EVs, especially their capacity to fuse with target cell membranes and deliver luminal cargo, is limited. Currently available EV-cargo delivery assays are indirect and thus unlikely to uncover molecular players and conditions that specifically control the EV-fusion step. Here, we present a novel live-cell imaging assay for detection of EV-binding, -uptake, and -fusion in time and space. We employed the SunTag system for exceptional signal amplification. EV-donor cells were engineered to tag the luminal EV-membrane with a fluorescent label coupled to SunTag peptides. Recipient cells express fluorescent single-chain anti-SunTag antibody (STAb), which binds EV-enclosed SunTag upon its cytosolic exposure. Using SunTagged EVs carrying fusogen VSV-G, we visualize the EV-fusion process, quantify fusion kinetics and efficiency, and determine subcellular localization of fusion events. We term this methodology *EV-FUSIM* (<u>E</u>xtracellular <u>V</u>esicle <u>Fu</u>sion <u>S</u>patiotemporal Imaging <u>M</u>ethod). In the future, this technology can support identification of fusogenic EV-subsets, as well as molecular players and drugs that modulate EV-fusion, without confounding effects of post-fusion processes. This will extend knowledge on EV-biology and can aid in the engineering of EVs that efficiently deliver intraluminal therapeutic payloads.

## Introduction

Extracellular Vesicles (EVs) are lipid-bilayer delimited particles of ∼50-500 nm released by virtually all cells across all kingdoms of life. They represent a heterogeneous mix of particles released through various biogenesis pathways. EVs contain a range of potentially functional molecules on their surface and inside their lumen, including proteins, nucleic acids and lipids (Van Niel et al., 2018; Buck & Nolte-’t Hoen, 2024; Welsh et al., 2024). In the past decade, EV-mediated communication has been shown to play a role in numerous (patho)physiological processes (Colombo et al., 2014; Kalluri & LeBleu, 2020). Additionally, EVs are regarded as enticing nano-vehicles for the delivery of therapeutic agents, based on their capacity to carry a range of cargos and their natural ability to modulate cellular processes (Elsharkasy et al., 2020; Herrmann et al., 2021; Cheng & Hill, 2022; Fusco et al., 2024).

Binding of EV-associated molecules to target cell receptors, either at the plasma membrane or at endosomal membranes following endocytosis and/or EV-membrane degradation, can trigger EV-induced signaling, e.g. during antigen presentation (Van Niel et al., 2018; Buzas, 2023; Hirosawa et al., 2025). EV-binding and -uptake have been extensively studied, for example by using fluorescently labelled or tagged EVs combined with flow cytometry or microscopy, revealing various underlying mechanisms (Toribio et al., 2019; Jackson Cullison et al., 2024; Hirosawa et al., 2025). However, there is a long-standing debate in the field on whether and how luminal EV-cargo contributes to cytosolic signaling (Somiya, 2020). Multiple mechanisms for EV-cargo delivery to the cytosol have been proposed. Most studies argue that EVs are taken up and deliver their cargo through fusion with endosomal limiting membranes, but fusion of EVs with plasma membranes has also been suggested (Parolini et al., 2009; Montecalvo et al., 2012). Besides fusion, cargo delivery may also occur through pore formation (Soares et al., 2015; Somiya, 2020). Currently available methods for detecting cargo delivery in human and mouse target cells are largely dependent on the functional activity of transferred cargo molecules, e.g. mRNA, miRNA, Cre-LoxP, CRISPR-Cas9 or split-enzyme reporters (Hung & Leonard, 2016; Lai et al., 2015; Zomer et al., 2016; De Jong et al., 2020; Bonsergent et al., 2021; Somiya & Kuroda, 2021a, 2021b; Bittel et al., 2021). Using these methods, evidence for cytoplasmic EV-mediated cargo delivery was found both *in vitro* and *in vivo*. Yet, the reported efficiency with which internalized EVs deliver functional cargo to the cytosol is generally low. For this reason, EVs designed for therapeutic cargo delivery are often equipped with viral fusion proteins, such as the vesicular stomatitis virus glycoprotein (VSV-G), to enhance their fusogenic properties (Zhang et al., 2020; Somiya & Kuroda, 2021a, 2021b; Bui et al., 2023; Ilahibaks et al., 2023; Ma et al., 2024; Liang et al., 2025; Obuchi et al., 2025). A downside of this approach is that virus-derived proteins can elicit immune reactions, which could preclude repeated dosing (Munis et al., 2019, 2020). The strong interest in optimizing current and developing novel strategies for EV-mediated therapeutic cargo delivery urges the need to increase our knowledge on how EVs are internalized and breach cellular membrane barriers to release luminal cargo into the cytoplasm.

Currently available reporter assays can provide evidence for EV-cargo delivery and illuminate parts of the underlying mechanisms, but only provide limited information on the actual EV-fusion event. Most reporter assays depend on activity-based read-outs, such as reporter activation upon genome editing by a delivered Cre-recombinase or production of bioluminescence by a reconstituted split luciferase enzyme (Zomer et al., 2016; Bonsergent et al., 2021), and are thus temporally disconnected from the actual EV-fusion event. This precludes acquisition of information on the real-time kinetics of this process. Furthermore, the reporter activity is determined by a cumulative series of events, including EV-binding, -uptake, -fusion and the use of cargo for a functional response. Moreover, these read-outs may be sensitive to inefficient enzyme complementation, cargo degradation or aberrant cargo localization, and tend to be spatially disconnected from the actual subcellular site of fusion. These limitations preclude the use of these reporter assays to study the efficiency and localization of discrete steps in the cascade of EV-binding, -uptake, -cargo delivery and signaling. Attempts have been made to directly visualize exposure of luminal EV-cargo to the cytosol of target cells using fluorescent cargo-targeting nanobodies (Joshi et al., 2020). However, this strategy only worked in cells after chemical fixation. This restricts analysis to a limited snapshot of the delivery events taking place over time and risks fixation-induced effects on the integrity of the endosomal system (Murk et al., 2003; Tanaka et al., 2010; Schnell et al., 2012). Live-cell imaging of the EV-fusion process represents an enticing alternative for visualization of EV-fusion in time and space. However, visualization of small amounts of cargo molecules exposed to the cytosol upon EV-fusion with limited laser exposure to safeguard cell viability requires exceptionally high detection sensitivity.

Here, we describe the development of a novel EV-binding, -uptake and -fusion reporter assay using state-of-the-art molecular tools and live-cell confocal microscopy (**Fig. 1**). For detection of EV-fusion events in live cells, we adapted the SunTag system, which was originally developed for live-cell single-molecule microscopy (Tanenbaum et al., 2014) and previously used to visualize highly dynamic processes such as cellular or viral protein translation (Yan et al., 2016; Boersma et al., 2020). In order to apply this system to study EV-cell interactions, we tagged the luminal leaflet of the EV-membrane with a red fluorescent protein coupled to SunTag peptides. EV-recipient reporter cells were engineered to express green fluorescent anti-SunTag single-chain antibodies in their cytosol. Upon EV-fusion, SunTag peptides are exposed to the cytosol of recipient cells, resulting in a strong local concentration of anti-SunTag and thus a detectable increase in green fluorescence intensity, visible as green spots. Dual color live-cell imaging of EV-recipient cells after addition of EVs isolated from EV-donor cells allows for simultaneous visualization of EV-binding and -uptake, as well as EV-fusion, in time and space. We term this methodology EV-FUSIM (<u>E</u>xtracellular <u>V</u>esicle <u>Fu</u>sion <u>S</u>patiotemporal Imaging <u>M</u>ethod).

**Figure 1.**
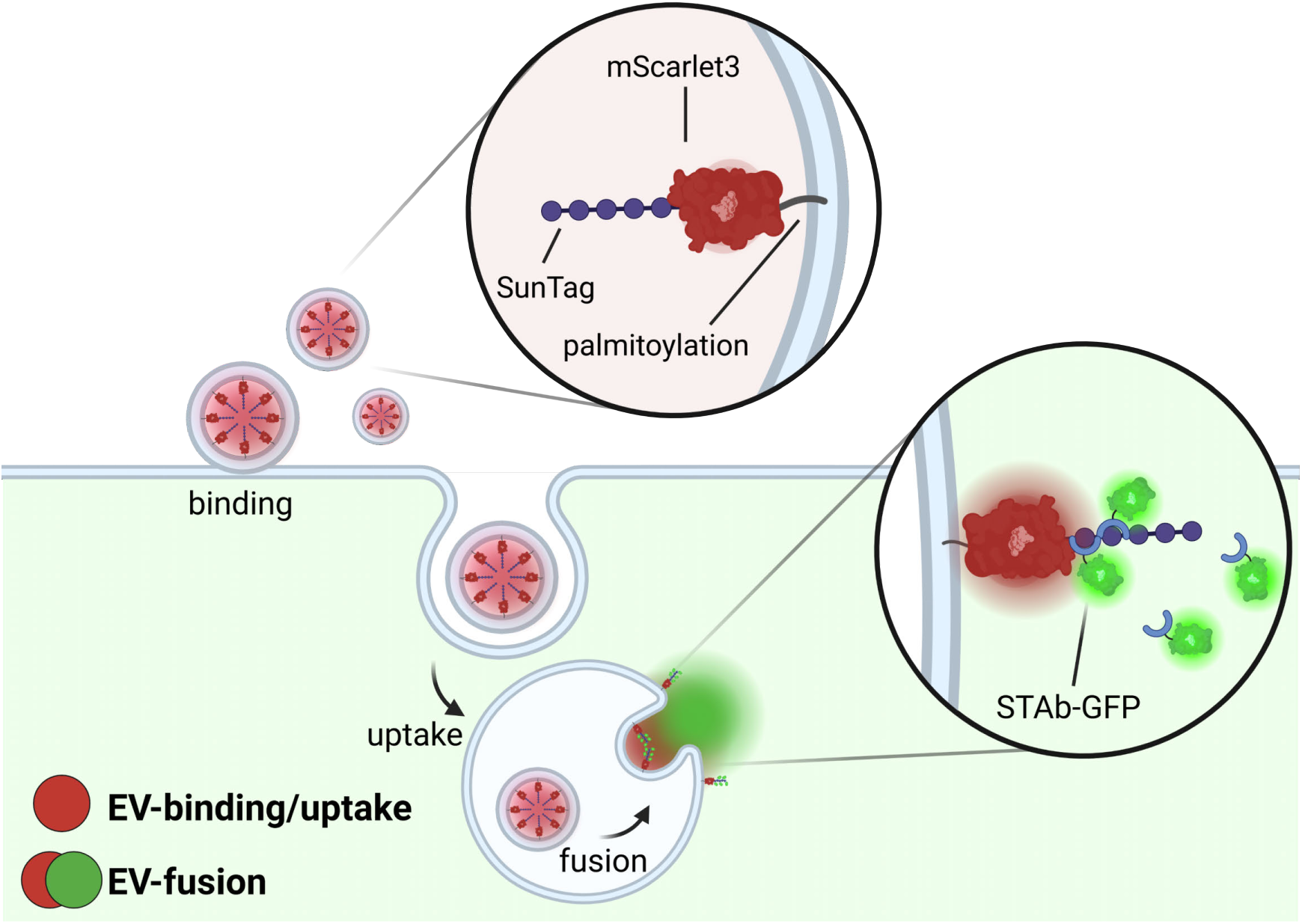
EV-FUSIM assay principle. EV-donor cells are engineered to release EVs containing luminal red fluorescence and SunTag, by lentivirally transducing them with a construct containing a palmitoylation motif, red fluorescent protein mScarlet3 and 10 repeats of the SunTag peptide. Upon addition to EV-recipient cells expressing green fluorescent anti-SunTag antibodies (STAb-GFP), EV-binding and -uptake will be visible as red fluorescent spots on and in the cells, whereas EV-fusion will result in exposure of luminal SunTag to the cytosol, thereby resulting in local concentration of STAb-GFP, visible as green spots. Created in BioRender. van den Ende, J. (2025) https://biorender.com/d7cuov7.

We first demonstrate that SunTag can be efficiently loaded with the proper orientation into HeLa cell-derived EVs. Proof-of-concept for the suitability of EV-FUSIM to visualize fusion events was obtained using HeLa cell-derived EVs mounted with VSV-G, which strongly increased fusion efficiency in HeLa EV-recipient cells. We demonstrate that simultaneous, real-time analysis of EV-binding, -uptake and -fusion provides the opportunity to uncouple effects of interference strategies on discrete steps in the EV-life cycle. Moreover, our data illustrates that EV-FUSIM can be used to explore the subcellular localization of EV-fusion events. This valuable addition to the EV research toolbox can assist in identification and characterization of fusogenic EV-subsets and fusion-promoting recipient cell states, as well as in optimization of cargo delivery by therapeutic (modified) EVs.

## Materials & Methods

### Cells

Human cervical carcinoma (HeLa R19) and human embryonic kidney (HEK293T) cells were purchased from the American Type Culture Collection (Rockville, MD; ATCC CCL-2 and ATCC CRL-3216, respectively). HeLa STAb cells, stably expressing AausFP1-STAb (denoted as STAb-GFP) and a blue fluorescent protein tagged with a nuclear localization signal (BFP-NLS), were derived from HeLa R19 as described previously (Tanenbaum et al., 2014; Boersma et al., 2020; Schipper et al., 2025) (STAb was originally derived from the plasmid pHR-scFv-GCN4-sfGFP-GB1-dWPRE, gift from Prof. Ron Vale – UCSF – also available as Addgene #60907). HeLa Gal3 cells, stably expressing mAG-Gal3 and BFP-NLS, were generated as described below. All cell lines were cultured in a humidified incubator at 37°C and 5% CO2 in Dulbecco’s Modified Eagle Medium (DMEM; high glucose, GlutaMAX^TM^, Thermo Fischer Scientific, Waltham, MA, USA), supplemented with 10% fetal bovine serum (FBS; Sigma-Alrich), 100 U/mL penicillin and 100 µg/mL streptomycin (Gibco, Paisley, United Kingdom).

### Plasmids

For luminal tagging of EVs in donor cells (palm-mScar3-10xSunTag), PCR-based cloning was used to insert a fusion construct containing a palmitoylation signal sequence, mScarlet3 fluorophore (mScar3), a short GS-linker (GGGGSGGGGS) and ten repeats of the SunTag peptide, into a pHAGE2 backbone. Insert donor plasmids were: pQC palm-tdTomato (gift from Dr. Tom Driedonks, made with commercial pQC backbone, Clontech, and insert originally derived from gift pCAG PalmtdTomato plasmid described by Prof. Xandra Breakefield) (Lai et al., 2015) for the palmitoylation sequence, pN1 p3xnls-mScarlet3 (gift from Prof. Dorus Gadella – Addgene #189775) (Gadella et al., 2023) for the mScarlet3 sequence and pcDNA4TO-UP-10xGCN4_v4-U2AF2 (generated in Tanenbaum lab) for the 10xSunTag (GCN4) sequence. For direct tagging of VSV-G, PCR-based cloning was used to insert mScar3, a GS-linker and ten repeats of the SunTag peptide, as described above, into a pCMV VSV-G plasmid (gift from Dr. Tom Driedonks, originally deposited on Addgene by Prof. Bob Weinberg - #8454) (Stewart et al., 2003). Palm-mScar3-10xSunTag and VSV-G-mScar3-10xSunTag plasmids will be made available to the community through Addgene upon publication, as #240626 and #240630, respectively.

For generation of endolysosomal rupture reporter cells (HeLa Gal3), a pHAGE2 mAG-Gal3 plasmid (gift from Prof. Niels Geijsen – Addgene #62734) (D’Astolfo et al., 2015) for expression of mAzamiGreen-tagged Galectin-3, and a previously described pHR NLS-BFP plasmid (Boersma et al., 2020) for tagging of the nucleus with Blue Fluorescent Protein, were used.

### Generation of stable transgenic HeLa cells

Lentiviruses were produced by co-transfection of HEK293T cells with a plasmid encoding the transgene of interest and packaging plasmids psPAX2 (gift from Dr. Tom Driedonks, originally deposited on Addgene by Prof. Didier Trono - #12260) and pCMV VSV-G into HEK293T cells using Lipofectamine 2000 (Thermo Fischer Scientific), according to the manufacturer’s protocol. Supernatants were harvested 3 days after transfection, subjected to 0.45 µm filtration and stored at −80°C until transduction. Lentivirus was supplemented with 8 µg/mL polybrene (Sigma-Aldrich) and added to HeLa R19 cells. Spin-infection was performed at 2000 rpm, 33°C for 90 minutes. After 3 days, transduced cells were selected using 5 µg/mL puromycin. HeLa Gal3 cells were additionally selected for high BFP and intermediate mAG fluorescence using flow cytometric sorting.

### Plasmid transfection

Plasmid transfection was performed at approximately 70% confluency using FuGENE HD transfection reagent (Promega) according to the manufacturer’s protocol, with a 3:1 reagent:DNA ratio and 80 ng (live-cell imaging in 96W format), 500 ng (immunofluorescence in 24 well format) or 15 µg (EV isolation in T75 flask format) plasmid DNA. After overnight transfection, experiments were conducted as described below for confluent, non-transfected cell layers.

### Antibodies

The following antibodies were used for western blotting and/or immunofluorescence staining: mouse-α-CD9 (1:2000, clone HI9a; Biolegend, San Diego, CA), mouse-α-CD63 (1:500, clone Ts63; Invitrogen, MA), mouse-α-GCN4 tag (1:2000, clone C11L34; Proteogenix, France), mouse-α-VSV-G (1:2000, clone P4D5; Sigma-Aldrich), mouse-α-actin (1:30,000, clone AC-15; Sigma-Aldrich), rabbit-α-Histone H3 (1:1000, polyclonal; Cell Signaling Technology, Danvers, MA), HRP-coupled goat-α-mouse (1:8000, polyclonal; Sigma-Aldrich), AF647-coupled goat-α-mouse (1:200, polyclonal; Abcam).

### Western blot

For analysis of cells, samples were lysed in RIPA buffer (40 mM Tris-HCl pH 8, 0.5% sodium deoxycholate, 1% Triton X-100, 150 mM sodium chloride, 0.1% sodium dodecyl sulfate) supplemented with protease inhibitor cocktail (Roche) and cleared by centrifugation at 15,000x*g* for 15 min. Protein concentration was determined using a Pierce BCA assay kit (Thermo Fischer Scientific) according to the manufacturer’s protocol. 5 µg protein from cell lysates was mixed with Laemmli sample buffer (LSB: 62.5 mM Tris-HCl pH 6.8, 2% SDS, 10% glycerol) with or without 20 mM 2-mercaptoethanol. For analysis of EVs within density gradient fractions, trichloroacetic acid (TCA) precipitation was performed to enable protein analysis. In brief, samples were supplemented with 125 µg/mL sodium deoxycholate and 10% ice-cold TCA, then pelleted for 15 min at 15,000xg. Pellets were washed 2x with ice-cold aceton and resuspended in LSB with or without 20 mM 2-mercaptoethanol. For analysis of purified EVs from ProtK protection experiments, samples were directly mixed with LSB. Proteins were denatured by incubating for 4 min at 98°C, separated on dodecyl sulfate-polyacrylamide gels by electrophoresis (SDS-PAGE), and transferred to 0.2 µm PVDF membranes using a Trans-Blot Turbo Transfer system (Bio-Rad). Membranes were blocked with 0.25% (v/v) fish skin gelatin (FSG; Sigma-Aldrich) in TBS + 0.1% Tween-20 (TBS-T) or 2% BSA in PBS + 0.1% Tween-20 (PBS-T) for 1h at RT, followed by incubation o/n with primary antibodies at 4°C. Membranes were washed 5x in the appropriate blocking buffer, followed by incubation for 45 min at RT with secondary antibodies. After 3x wash in PBS-T or TBS-T, membranes were developed using Clarity Western ECL solution (BioRad) for detection on an Amersham ImageQuant 800 Imager. Imageswere processed using the ImageQuantTL software package (v10.2, Cytiva).

For SunTag Western blot detection, two bands were observed. The top bands run near the height expected for the full-length palm-mScar3-10xSunTag construct (predicted to be 55.75 kDa). The bottom bands are likely the result of hydrolysis of the acylimine moiety in mScar3, as has been shown for dsRed-like fluorescent proteins during denaturation conditions required for Western blotting (Gross et al., 2000; McCullock et al., 2020). The lower band was relatively more prominent in TCA-precipitated samples (Fig. 2C, Fig. 3B) compared to non-precipitated samples (Fig. 2E, Fig. 3D), which is likely the result of strong protein denaturation during TCA precipitation (Koontz, 2014).

**Figure 2.**
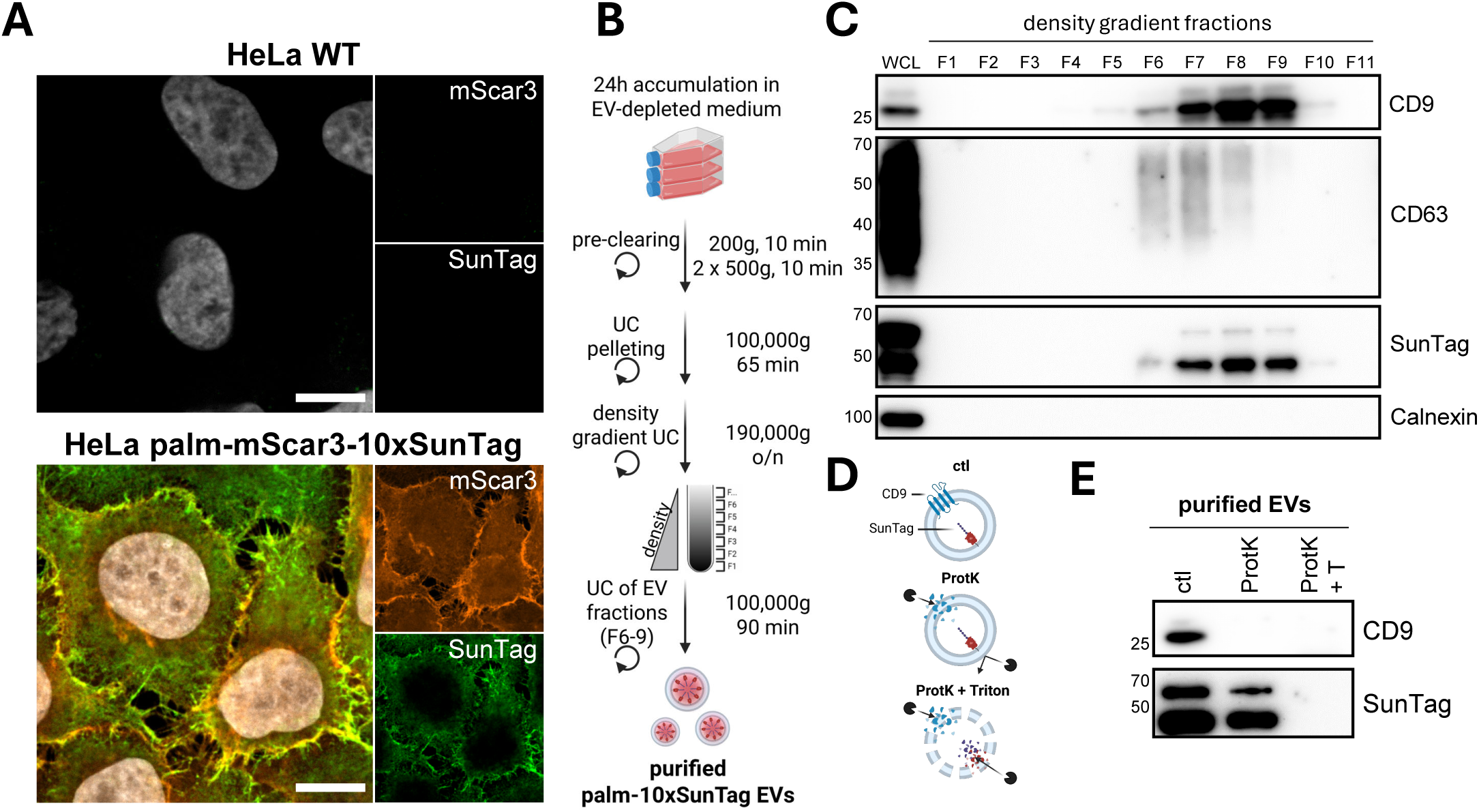
HeLa palm-mScar3-10xST cells release luminally SunTagged EVs. **(A)** HeLa WT and palm-mScar3-10xST cells fixed after 24h incubation in EV-depleted medium and immunolabeled for SunTag. Images shown are mid-nuclear Z-slices, showing fluorescence for mScar3 (red), SunTag (green) and Hoechst-labeled nuclei (white). Scale bars represent 10 µm. Images are representative of n=2 independent experiments. **(B)** Schematic overview of EV-isolation methodology. See Materials & Methods for detailed description. Created in BioRender. van den Ende, J. (2025) https://biorender.com/hvvdrpk. **(C)** EV-containing ultracentrifugation pellets from HeLa palm-mScar3-10xST cells were floated into iodixanol density gradients, after which individual gradient fractions were concentrated using TCA precipitation and analyzed for the presence of protein markers by Western blotting. For SunTag, two bands are visible, corresponding to full-length construct (top) and a fragmented version (bottom) which is likely the result of denaturation in the Western blot procedure (see Materials & Methods for detailed explanation). **(D)** Schematic overview of Proteinase K protection assay principle. See Materials & Methods for detailed description. Created in BioRender. van den Ende, J. (2025) https://BioRender.com/pb45em2. **(E)** Gradient-purified EVs were treated with ProtK in the absence or presence of 0.2% Triton X-100 and analyzed by Western blotting. For SunTag, two bands are visible, corresponding to full-length construct (top) and a fragmented version (bottom) which is likely the result of denaturation in the Western blot procedure (see Materials & Methods for detailed explanation).

**Figure 3.**
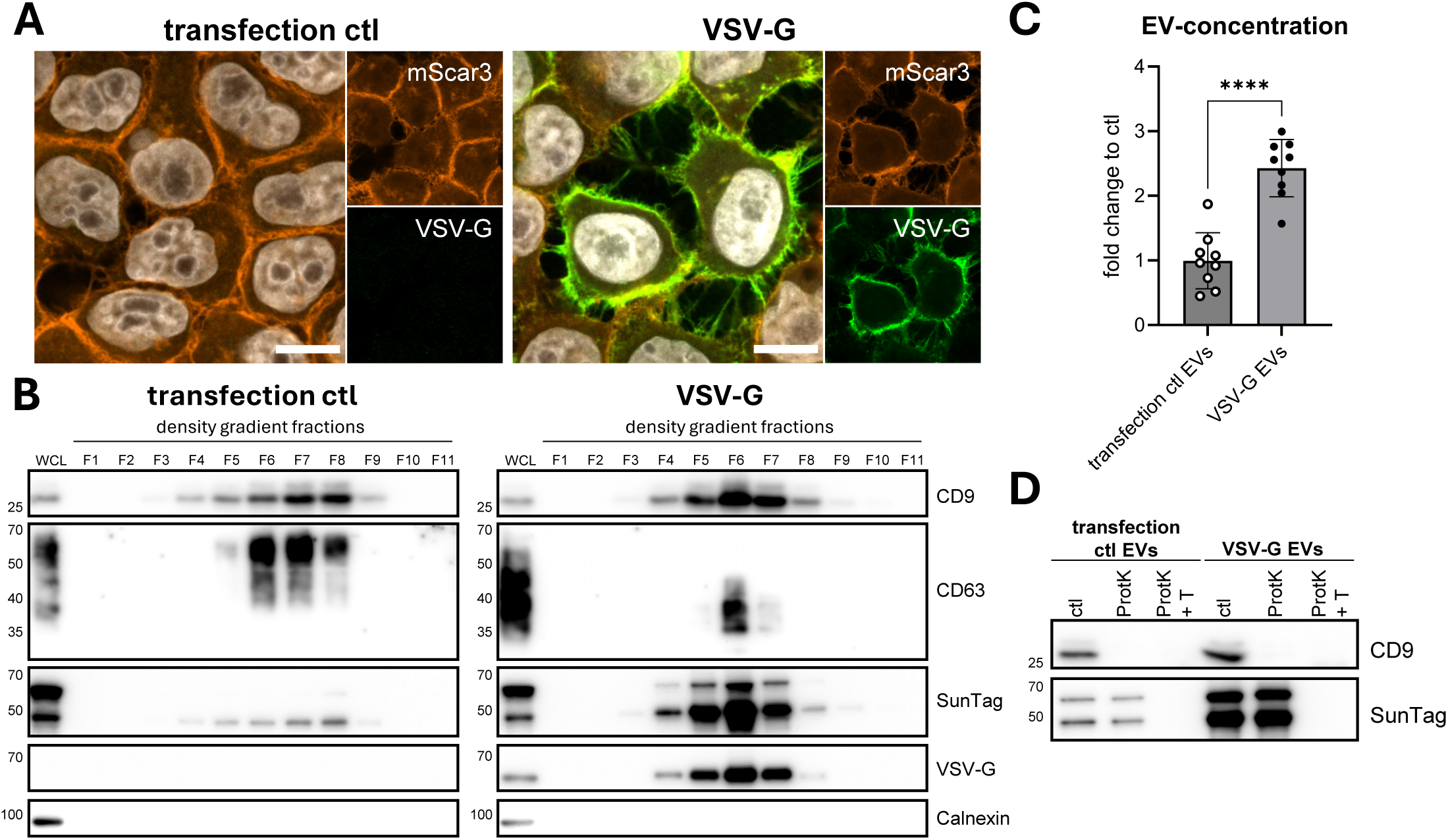
Transfection of HeLa palm-mScar3-10xST cells with VSV-G results in release of VSV-G-containing SunTagged EVs. **(A)** Mock- or VSV-G-transfected HeLa palm-mScar3-10xST cells were fixed after overnight transfection followed by 24h incubation in EV-depleted medium, then immunolabeled for VSV-G. Images shown are mid-nuclear Z-slices, showing fluorescence for mScar3 (red), VSV-G (green) and Hoechst-labeled nuclei (white). Scale bars represent 10 µm. Images are representative of n=2 independent experiments. **(B)** EV-containing ultracentrifugation pellets from mock- and VSV-G-transfected cells were floated into iodixanol density gradients, after which individual gradient fractions were concentrated using TCA precipitation and analyzed for the presence of protein markers by Western blotting. For SunTag, two bands are visible, corresponding to full-length construct (top) and a fragmented version (bottom) which is likely the result of denaturation in the Western blot procedure (see Materials & Methods for detailed explanation). **(C)** EV-concentrations in gradient-purified EV preparations isolated from mock- and VSV-G-transfected cells were quantified using high resolution flow cytometry. Graph shows the mean fold change to transfection ctl EVs ± SD, calculated from n=8 independent experiments. **** p≤0.0001 as determined by paired t-test. **(D)** Gradient-purified EVs isolated from mock- and VSV-G-transfected cells were treated with ProtK in the absence or presence of 0.2% Triton X-100 and analyzed by Western blotting. For SunTag, two bands are visible, corresponding to full-length construct (top) and a fragmented version (bottom) which is likely the result of denaturation in the Western blot procedure (see Materials & Methods for detailed explanation).

### Immunofluorescence staining

After 24h incubation with EV-depleted medium, cells grown on coverslips were washed and then fixed with 4% PFA for 15 min, washed 2x with PBS + 10 mM glycine, 1x with PBS and permeabilized for 10 min with 0.2% Triton X-100. After washing 3x with PBS + 0.1% Tween-20 (PBS-T), cells were blocked with 2% BSA in PBS-T for 1h at RT, then stained for 45 min with primary antibodies diluted in block buffer. After three washes with PBS-T, antibody labelling was repeated with fluorophore-conjugated secondary antibodies and Hoechst 33342 (1 µg/mL, Thermo Fischer Scientific) for 30 min at RT. Finally, coverslips were washed with PBS-T and MQ before being mounted in ProLong Diamond Antifade Mountant (Thermo Fischer Scientific). Mounted slides were imaged according to specifications described below.

### EV isolation

Confluent cell layers were washed three times with PBS +Ca +Mg and incubated with cell culture medium containing 10% EV-depleted FBS. To generate this EV-depleted FBS, FBS was prediluted 1:3 in DMEM, ultracentrifuged for 16-20h at 28,000 rpm (100,000x*g*) in an SW32 rotor (k-factor 256.8) and passed through a 0.22 µm filter. 24h after replacing the medium, cell culture supernatants were harvested and centrifuged for 10 min at 200x*g* and 2x 10 min at 500x*g* to remove cells and cell debris. Cleared supernatant samples were treated with 5 mM MgCl2 and 0.1 mg/mL DNase I (Roche) for 30 min at 37°C, after which they were supplemented with 25 mM HEPES (Sigma-Aldrich). EVs were enriched from clarified supernatants by UC pelleting for 65 min at 28,000 rpm (100,000x*g*) in an SW32 rotor (k-factor 256.8) and resuspended PBS +0.1% BSA (cleared from aggregates by ultracentrifugation for 16-20h at 28,000 rpm - 100,000x*g*). Next, EVs were mixed with iodixanol (Optiprep; Axis-Shield, Oslo, Norway) to a final concentration of 45% and overlaid with a linear gradient of 40-5% iodixanol in PBS by adding 8 layers of 1.3 mL, decreasing in density at 5% decrements. Density gradients were centrifuged at 39,400 rpm (192,000x*g*) for 16h in a SW41 rotor (k-factor 124). Gradient fractions of 1 mL were collected from the top and their densities assessed by refractometry. For experiments with purified EVs, gradient fractions 6-9 were diluted in 30 mL PBS +0.1% BSA (cleared from aggregates), isolated by UC pelleting for 90 min at 28,000 rpm in an SW32 rotor (k-factor 256.8) and resuspended in PBS.

We have submitted all relevant data regarding methodological reporting of our experiments to the EV-TRACK knowledgebase (EV-TRACK ID: EV250079) (EV-TRACK Consortium et al., 2017).

### Proteinase K digestion

Purified EVs, isolated by UC pelleting from Optiprep density gradient fractions, were resuspended in PBS and incubated in the absence or presence of 0.2% Triton X-100 for 5 min at RT. Then, samples were incubated in the absence or presence of 45.45 ng/µL Proteinase K (Roche, Basel, Switzerland) for 1h at 37°C. Proteinase K was inactivated by the addition of 2 mM PMSF (Roche) and subsequent incubation for 15 min on ice. Samples were mixed with LSB for SDS-PAGE and Western blot analysis.

### High resolution flow cytometric analysis of EVs

For assessment of EV distribution in Optiprep density gradients, EVs were fluorescently labeled prior to density gradient centrifugation with 30 µM CFSE (Invitrogen, Carlsbad, CA) for 60 min at RT. Individual gradient fractions 4-11 were diluted 1:20 in PBS immediately prior to high-resolution flow cytometric analysis on a Cytek Aurora spectral flow cytometer, using optimized settings for the detection of small particles, with a fluorescent threshold of 1,050 in the B2 detector. A fixed volume of 20 uL of sample was measured to allow for quantitative comparison of EV release.

For normalization of purified EVs for live-cell imaging experiments, EVs isolated by UC pelleting from Optiprep density gradient fractions were resuspended in PBS. Samples were further diluted 1:400 in PBS immediately prior flow cytometric analysis, using similar settings as described above but with a SSC threshold of 2,000. A fixed volume of 5 µL of samples was measured.

Data analysis for flow cytometric experiments was performed using Spectroflo v3.3.0. Background events detected in PBS were subtracted and EV numbers in each sample were calculated based on their respective dilution factors.

Relevant data regarding methodological reporting of flow cytometry experiments has been attached as **Supplementary Tables 2 and 3** in the form of completed MIFlowCyt and MIFlowCyt-EV checklists, respectively (Lee et al., 2008; Welsh et al., 2020).

### Live-cell imaging of EV-recipient cells

CELLview imaging chambers (75×25mm, glass bottom; Greiner Bio-One, The Netherlands) were seeded with EV-recipient cells one day prior to imaging in order to be confluent the next day, in 150 uL DMEM without phenol red (Thermo Fischer Scientific), supplemented with 10% FBS (Sigma-Alrich), 100 U/mL penicillin and 100 µg/mL streptomycin (Gibco). After installation on the microscope and potential drug or dye pre-treatments as described below, 100 uL medium was removed and replaced with the desired concentration of EVs pre-diluted in medium to a total volume of 100 uL, after which time-lapse imaging was immediately started as described below.

#### Drug treatments

To investigate the dependence of VSV-G-mediated EV-fusion on endolysosomal acidification, EV-recipient cells were pre-treated with 200 nM bafilomycin A1 (Cayman Chemical, Ann Arbor, MI) or DMSO vehicle control for 30 min prior to EV-addition, after which EVs were added as described above in the presence of the same concentrations of drugs. To induce endolysosomal rupture, recipient cells were treated with 1 mM L-Leucyl-L-Leucine methyl ester (hydrochloride) (LLOME; Cayman Chemical) immediately prior to live-cell imaging.

#### Dye staining

To investigate localization of EV-fusion events at endocytic compartments, EV-recipient cells were incubated with 20 nM LysoTracker-DeepRed or 5 µg/mL CellMask-DeepRed (both Invitrogen) for 15 min. After incubation, staining solution was removed and cells were washed once with medium, following EV-addition as described above.

### Microscopical image acquisition

Images were acquired using an Olympus SpinSR10 spinning disk confocal microscope (Evident, Leiderdorp, The Netherlands) equipped with an ORCA Fusion sCMOS camera (Hamatsu, Japan) and a 100X oil immersion objective (UPLXAPO 100XO, NA1.45; Olympus). Image recording was operated with CellSens Dimension software (v4.2) using a quadband main dichroic mirror (D405/488/561/640nm). Hoechst/NLS-BFP were detected after 405 nm excitation using a B477/60 nm emission filter. STAb-GFP/mAzamiGreen-Gal3 were detected after 488 nm excitation using a B515/30 nm emission filter. EV-mScar3 was detected after 561 nm excitation using a B607/36 nm emission filter. AF647/LysoTracker-DeepRed/CellMask-DeepRed were detected after 640 nm excitation using a B685/40 nm emission filter.

Live-cell image acquisition post-EV-addition.

For all experiments, Z-stacks were recorded at several randomly pre-selected positions throughout the wells of interest (minimum of three), at the time interval designated in the appropriate figure legends. For STAb and Gal3 experiments, 11 Z-slices of 1 µm step-size were recorded. For STAb localization experiments, 21 Z-slices of 0.5 µm step-size were acquired.

### Analysis of live-cell imaging experiments

Snapshot images (single Z-slices or maximum intensity projections, as denoted in figure legends) were exported from CellSens Dimension software (v4.2).

For quantitative analysis, image pre-processing was performed in FIJI (v1.54p) (Schindelin et al., 2012) to remove single pixel intensity outliers and to set the intensity range of all channels as twice the minimal intensity value in the field-of-view as minimum and 5000 as maximum. Images were median-filtered (radius = 2) and exported to a native readable h5-Imaris file format using the BigDataProcesser2 plug-in (Tischer et al., 2021). Imaris software (vs10.2; Oxford Instruments, UK) was used for further analysis in 3D, including cell segmentation and spot detection (scripts and parameters used for object creation can be found in **Suppl. Table 1**). In brief, cells were segmented using the cell-associated STAb-GFP or mAG-Gal3 channel by creating Surface objects using the machine-learning module. Strongly bright cells were filtered out from processing to reduce false positive STAb/Gal3 spot detection. Subsequently, EV-mScar3 and STAb spots were segmented using the Spot detection module. Spots were filtered based on the distance to the created cell Surface objects to exclude non-cell-associated spots. For EV-mScar3 spots, an additional classification was performed based on the distance to the cell Surface to differentiate plasma membrane-proximal and -distal spots, in order to differentially quantify EV-binding and -uptake. Object counts for STAb or Gal3 spots, EV-mScar3 total spots, EV-mScar3 PM-proximal spots and EV-mScar3 PM-distal spots, as well as the sum volume of cell objects, were exported per timepoint for each field-of-view and further analyzed in Excel (v2408; Microsoft, US). Average sum cell volumes/field-of-view were calculated for both STAb and Gal3 datasets (132290.49 and 111537.98 µm^3^/FOV, respectively), after which all spot counts were corrected for the difference between the associated sum cell volume and the calculated average, in order to correct for differences in cell coverage and occasional focus losses. Maximum corrected spot counts within the full timelapse for each field-of-view were used for bar graphs and statistical comparison between conditions.

#### Localization analyses

For LysoTracker- and CellMask-DeepRed channels, images were subjected to additional background subtraction (radius = 0.3 µm) in Imaris, after which Surface objects were segmented in these channels using absolute intensity thresholding. For NLS-BFP channels, Surface objects were created using the machine-learning module. For all three, binary masks were generated from these Surface objects to use in classification of overlapping STAb spots. Cell Surface objects and STAb spots were segmented as described above. As a threshold for overlap, we chose a minimum of 25% overlap of STAb spot and localization binary masks. We determined the average STAb spot object volume in the localization dataset to be 1.96 µm^3^, corresponding to 928 voxels (with dimensions of 0.065 * 0.065 * 0.5 µm each). Thus, a minimum of 25% overlap corresponds to 232 voxels. As each positive voxel in the binary masks has an intensity of 1, a summed intensity of 232 in the masked channel was used as threshold to designate a spot as an overlapping event. Average sum cell volumes/field-of-view were calculated for the STAb localization dataset (122418.73 µm^3^/FOV), and used for correction of spot counts as described above.

### Statistical analysis

Statistical analysis and data plotting was performed using Prism (10.0; GraphPad Software). Statistical tests used are denoted in the appropriate figure legends.

### Table-of-contents graphical abstract

Created in BioRender. van den Ende, J. (2025) https://biorender.com/8ihhtg1.

## Results

### Engineering of donor cells for production of SunTagged EVs

In order to simultaneously detect EV-binding/uptake and -fusion upon addition to recipient cells, we developed a strategy to tag the lumen of EVs released by mammalian cells with a red fluorescent protein and SunTag peptides (**Fig. 1**). By implementing a repeating array of SunTag peptides, small amounts of cargo molecules exposed to the cytosol upon EV-fusion can be detected by multiple fluorescently labelled single-chain anti-SunTag antibodies (STAb-GFP) per tagged molecule, facilitating strong signal amplification. To achieve this, we generated HeLa EV-donor cells stably expressing mScarlet3 and 10 repeats of SunTag linked to a palmitoylation motif (HeLa palm-mScar3-10xST), a proven strategy for tagging the inner leaflet of the EV-membrane (Lai et al., 2015). Microscopic imaging of these cells confirmed expression and colocalization of mScarlet3 and SunTag, predominantly at the plasma membrane (**Fig. 2A**). EVs released by these cells were isolated from cell culture supernatants using differential (ultra)centrifugation followed by density gradient centrifugation (**Fig. 2B**). Immunoblot analysis confirmed that SunTag co-fractionated with EV-associated tetraspanins CD9 and CD63 in fractions 6-9 of the density gradient (**Fig. 2C**). These fractions, with densities of 1.04 – 1.10 g/mL, contained the highest number of EVs as assessed by high resolution flow cytometry (**Suppl. Fig. 1A-C**). To prove the luminal orientation of EV-associated SunTag, we tested the pooled EV fractions (1.04 – 1.10 g/mL) for protection of SunTag from Proteinase K (ProtK), which digests externally accessible EV proteins but leaves those protected by the EV-membrane intact (**Fig. 2D**). Indeed, ProtK treatment did not lead to SunTag digestion unless the EV-membrane was disrupted with the detergent Triton X-100, while CD9 was also cleaved by ProtK in the absence of detergent (**Fig. 2E**). Based on these data, we conclude that engineered EV-donor cells release EV-associated SunTag with the correct membrane orientation.

Incorporation of fusogenic viral glycoprotein VSV-G is a proven tool for enhancing cytosolic delivery of luminal EV-cargo (Zhang et al., 2020; Somiya & Kuroda, 2021a, 2021b; Ilahibaks et al., 2023; Bui et al., 2023; Ma et al., 2024; Liang et al., 2025; Obuchi et al., 2025). We incorporated VSV-G in our system in order to generate EVs with high fusogenic potential. Transfection of palm-mScar3-10xST EV-donor cells with VSV-G resulted in colocalization of palm-mScar3-10xST and VSV-G at the plasma membrane (**Fig. 3A**). Released VSV-G co-fractionated with CD9, CD63 and SunTag (**Fig. 3B**). VSV-G transfection led to a shift of EVs containing CD9/63 and SunTag to slightly higher densities (1.08-1.13 g/mL) when compared to EVs from control transfected cells (**Fig. 3B**). Additionally, CD63 was detected at a lower molecular weight in both cell lysates and EVs from VSV-G-transfected cells, suggesting altered glycosylation. In line with previous reports (Bui et al., 2023; Ilahibaks et al., 2023), VSV-G transfection led to a strong increase in total EV release, as measured with high resolution flow cytometry (**Fig. 3C, Suppl. Fig. 2**), and by immunoblotting for CD9 and SunTag (**Fig. 3D**). VSV-G transfection did not alter the orientation of EV-associated palm-mScar3-10xST, as demonstrated by ProtK protection assays (**Fig. 3D**). Together, these data demonstrate that transfection of EV-donor cells with VSV-G induces the release of VSV-G-containing SunTagged EVs. We used these EV-isolates to further develop our EV-FUSIM live-cell imaging assay.

### EV-FUSIM simultaneously traces EV-binding, -uptake and -fusion in live cells

EV-recipient cells were engineered to express cytosolic anti-SunTag single chain antibodies coupled to a bright green fluorescent protein (AausFP1 (Lambert et al., 2020), denoted as STAb-GFP from here onwards). EV-FUSIM requires that cytosolic exposure of SunTag upon EV-fusion locally concentrates STAb-GFP in recipient cells, which can be detected as fluorescent spots using live cell imaging. Direct transfection of HeLa cells stably expressing STAb-GFP with the SunTag construct, thereby bypassing the EV-fusion process, confirmed recruitment of STAb-GFP to mScar3-positive sites at the plasma membrane (**Suppl. Fig. 3**). Next, we tested the suitability of this system for live-cell imaging of EV-binding/uptake and -fusion (**Fig. 4A**). EVs were purified from HeLa palm-mScar3-10xST donor cells as described before and quantified using high resolution flow cytometry. Purified EVs were added to HeLa STAb-GFP recipient cells after which timelapse imaging was immediately initiated. Images were acquired from multiple positions per well, capturing Z-stacks with red- and green-channel fluorescence from each position at a time interval of 1 hour. Here, we treated 4 * 10^4^ recipient cells with a low (1 * 10^8^, 2500/cell) or high (5 * 10^8^, 12500/cell) dose of EVs, as well as a low dose of transfection control or VSV-G EVs. In all EV-treated conditions but not the medium control, cell-associated red fluorescent (mScar3+) spots accumulated over time, indicating binding and/or uptake of EVs (**Fig. 4B, Suppl. Fig. 4A**). Only for VSV-G EVs, this was also associated with an accumulation of green fluorescent (GFP+) spots, indicating EV-fusion, predominantly at early timepoints (**Fig. 4B, Suppl. Fig. 4B**).

**Figure 4.**
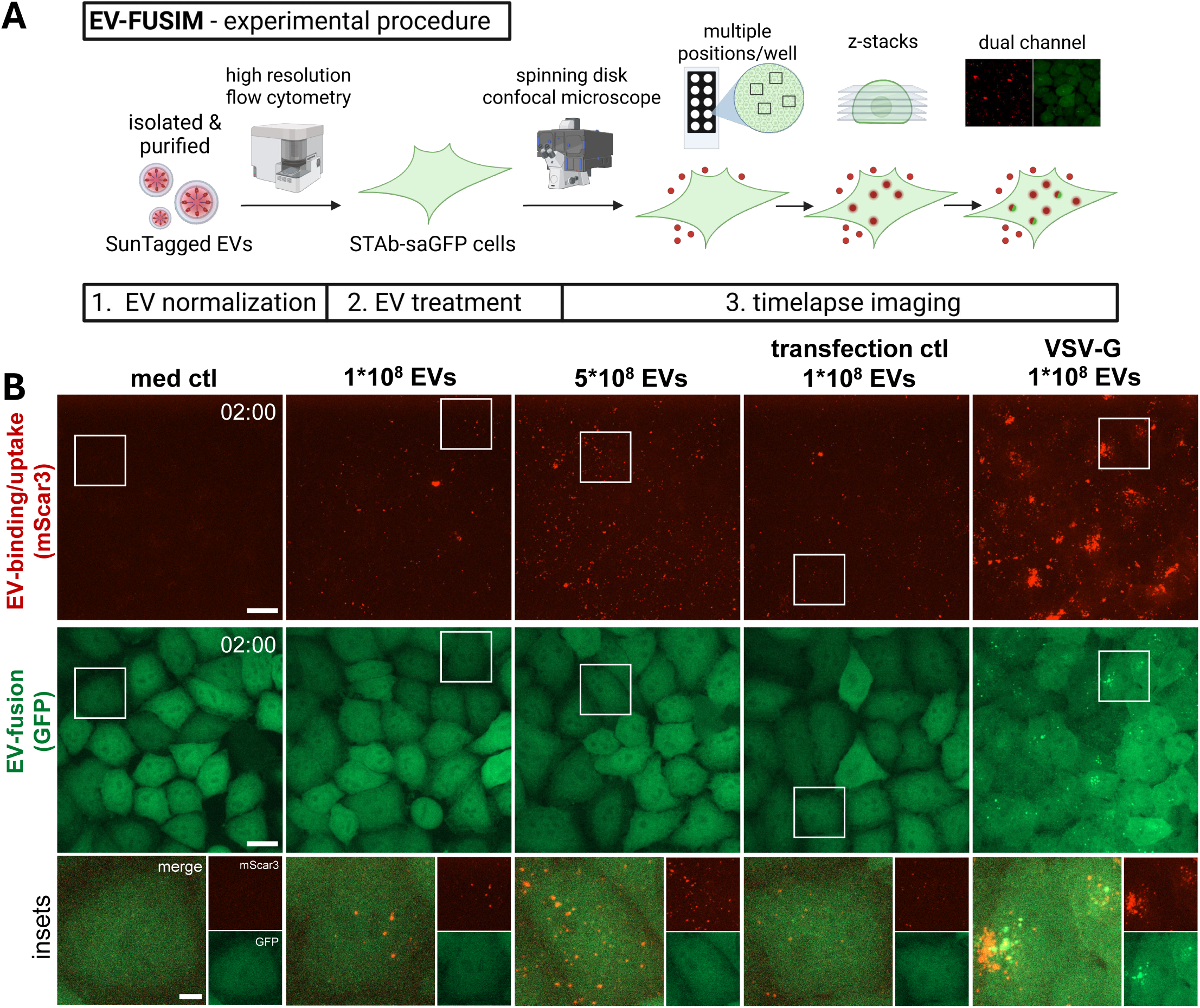
Live-cell imaging of HeLa STAb-GFP cells after addition of SunTagged EVs allows for tracing of EV-binding/uptake and -fusion over time. **(A)** Schematic overview of live-cell imaging procedure. See Materials & Methods for detailed description. Created in BioRender. van den Ende, J. (2025) https://BioRender.com/gaf5kzg. **(B)** Live HeLa STAb-GFP cells were subjected to timelapse imaging immediately after addition of medium only or medium containing the indicated concentrations and types of EVs, taking Z-stacks at a 1h time interval. Images are maximum intensity projections acquired at 2h post-EV-addition, showing fluorescence channels for mScar3 **(top)** and GFP **(middle)** in the same fields-of-view. Gamma was adjusted to 2 (mScar3) or 1.2 (GFP) for visualization purposes only. Scale bars represent 20 µm. White insets correspond to magnifications (**bottom**), which show mScar3, GFP and merged channels. Scale bar represents 5 µm. Images are representative of n=3-4 independent experiments.

For in-depth quantitative analysis of EV-binding/uptake and -fusion, we designed a 3D image analysis pipeline (**Fig. 5A**). Following image pre-processing, cell boundaries were detected and segmented in 3D, after which cell-associated fluorescent spots were detected in both channels and counted (see Materials & Methods for detailed description). Incubation of cells with EVs led to time- and dose-dependent accumulation of EV-binding/uptake (**Fig. 5B, C**). For VSV-G EVs, the EV-binding/uptake efficiency was strongly increased. Quantitative image analysis confirmed that EV-fusion was detected exclusively for VSV-G EVs (**Fig. 5D, E**). For EVs lacking VSV-G, neither a higher dose nor prolonged incubation significantly increased the detection of green fluorescent spots compared to the medium control. Both cell surface-bound and internalized EVs could be detected in these experiments, ruling out that a lack of EV-fusion detection is due to a lack of EV-uptake into the endosomal system (**Suppl. Fig. 5**). For VSV-G EVs, green fluorescent spots were detected immediately after the start of imaging and increased in number up to the first hour after addition of EVs (**Fig. 5D, E**). Beyond this timepoint, the detection of fusion events plateaued and eventually dissipated. Further corroborating efficient exposure of SunTag to the cytosol from VSV-G-containing palm-SunTag EVs, we observed a relative depletion of STAb-GFP signal from the nucleus, which is a known proxy for STAb sequestration in the cytosol (**Suppl. Fig. 6A-C**) (Khuperkar et al., 2020). At later timepoints, we observed an increase of plasma membrane-associated fluorescence, which precluded confident spot detection beyond 4 hours after EV-addition (**Suppl. Fig. 4B**). The observed disappearance of STAb concentrates from EV-fusion locations and redistribution to the plasma membrane is likely due to de-palmitoylation which frequently occurs in the cytosol (Kathayat & Dickinson, 2019), and may be followed by re-palmityolation and insertion in the plasma membrane (**Suppl. Fig. 6D, E**). Thus, EV-FUSIM readily detects EV-binding, -uptake and VSV-G-mediated EV-fusion in live cells over prolonged periods of imaging, allowing for quantitative assessment of these processes.

**Figure 5.**
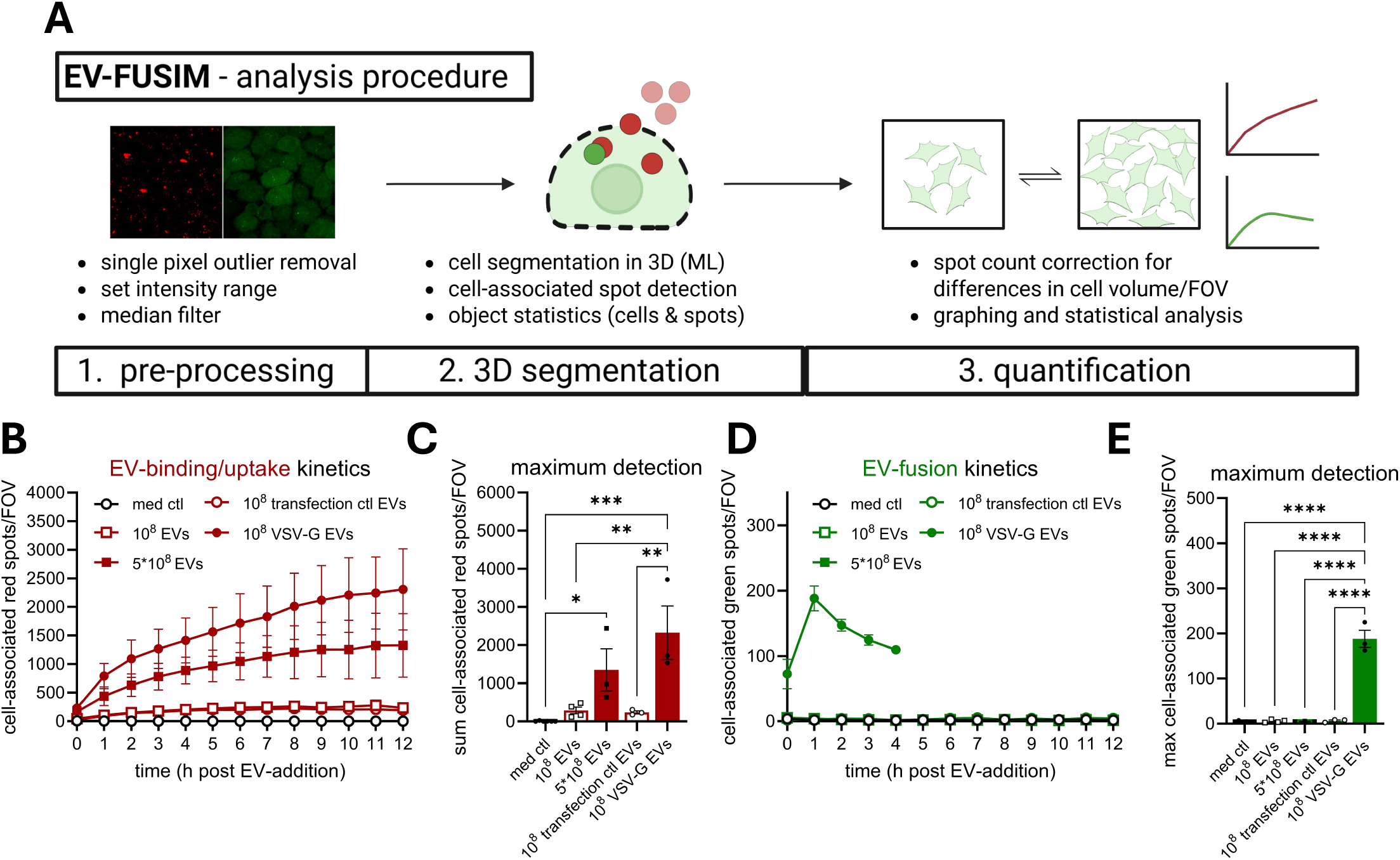
Image analysis with 3D cell segmentation facilitates quantitative assessment of EV-binding/uptake and -fusion. **(A)** Schematic overview of image analysis procedure. See Materials & Methods for detailed description. Created in BioRender. van den Ende, J. (2025) https://BioRender.com/k3ijxlv. **(B-E)** After segmentation of cells in 3D based on the GFP channel, cell-associated fluorescent spots in the red and green channel were counted for each field-of-view. In tandem, the total cell volume per field-of-view was counted, which was used to correct spot counts for differences to the average cell volume per field-of-view. Graphs show the corrected mean spots per timepoint **(B, D)** or mean maximum spot detection over the course of the experiment **(C, E)** per field-of-view for both mScar3 and GFP channels ± SEM, calculated from n=3-4 independent experiments with 3-6 fields-of-view per condition each. For all experimental conditions quantification was performed for all timepoints, except for GFP spots in the VSV-G EV condition, where only the first 4h were processed as changes in membrane fluorescence after this timepoint precluded proper spot detection. * p≤0.05, ** p≤0.01, *** p≤0.001 and **** p≤0.0001 as determined by one-way ANOVA with Tukey’s multiple comparisons test.

### Real-time visualization of EV-fusion events

The data in **Fig. 5** indicate that VSV-G-mediated EV-fusion has fast kinetics, peaking as early as 1 hour post-EV-addition. Therefore, we next used EV-FUSIM to assess EV-binding/uptake and -fusion kinetics at higher temporal resolution, by imaging cells every 15 min up to 2 hours after EV-addition (**Suppl. Video 1**). Even at these early timepoints, cell-associated red spots accumulated much more rapidly in cells treated with VSV-G EVs compared to control EVs (**Fig. 6A-C**). In cells treated with VSV-G-EVs but not control EVs, green spot detection increased from 15 to 60 min after addition of EVs, after which a plateau was reached (**Fig. 6A, D, E**). Thus, EV-FUSIM allows for assessment of fusion kinetics. Furthermore, imaging at a 1 min interval allowed for visualization of EV-fusion from individual endosomes in real-time, with red-fluorescent EV-containing endosomes eventually turning red-green positive upon EV-fusion (**Fig. 6F**). To exclude that the observed STAb-GFP concentrates were the result of endo-lysosomal rupture events rather than fusion of the EV- and endosomal membranes, we used an established Galectin-3 reporter assay in parallel live-cell imaging experiments (**Suppl. Fig. 7A**). Galectin-3 is a small cytosolic molecule which clusters on permeabilized endosomes by binding β-galactosides present on the inner leaflet of the endosomal membrane (Paz et al., 2010; D’Astolfo et al., 2015). HeLa cells were transduced to express Galectin 3 tagged with green fluorophore mAzamiGreen (HeLa mAG-Gal3) and used as EV-recipient cells in our live-cell imaging setup. Treatment with VSV-G EVs did not lead to an increase in green spots compared to untreated cells despite efficient EV-binding/uptake by the HeLa mAG-Gal3 reporter cells (**Fig. 6G-I, Suppl. Fig. 7B-D**). As a positive control, we used L-leucyl-L-leucine methyl ester (LLOMe). This polymer, described to permeabilize endo-lysosomes, rapidly induced accumulation of intracellular green spots (**Fig. 6G-I**). Thus, VSV-G-mediated EV-cargo delivery occurs through fusion of EV- and endosomal membranes, rather than endo-lysosomal rupture. Taken together, these data show that EV-FUSIM facilitates real-time, simultaneous assessment of EV-binding/uptake and -fusion of VSV-G EVs with unprecedented temporal resolution.

**Figure 6.**
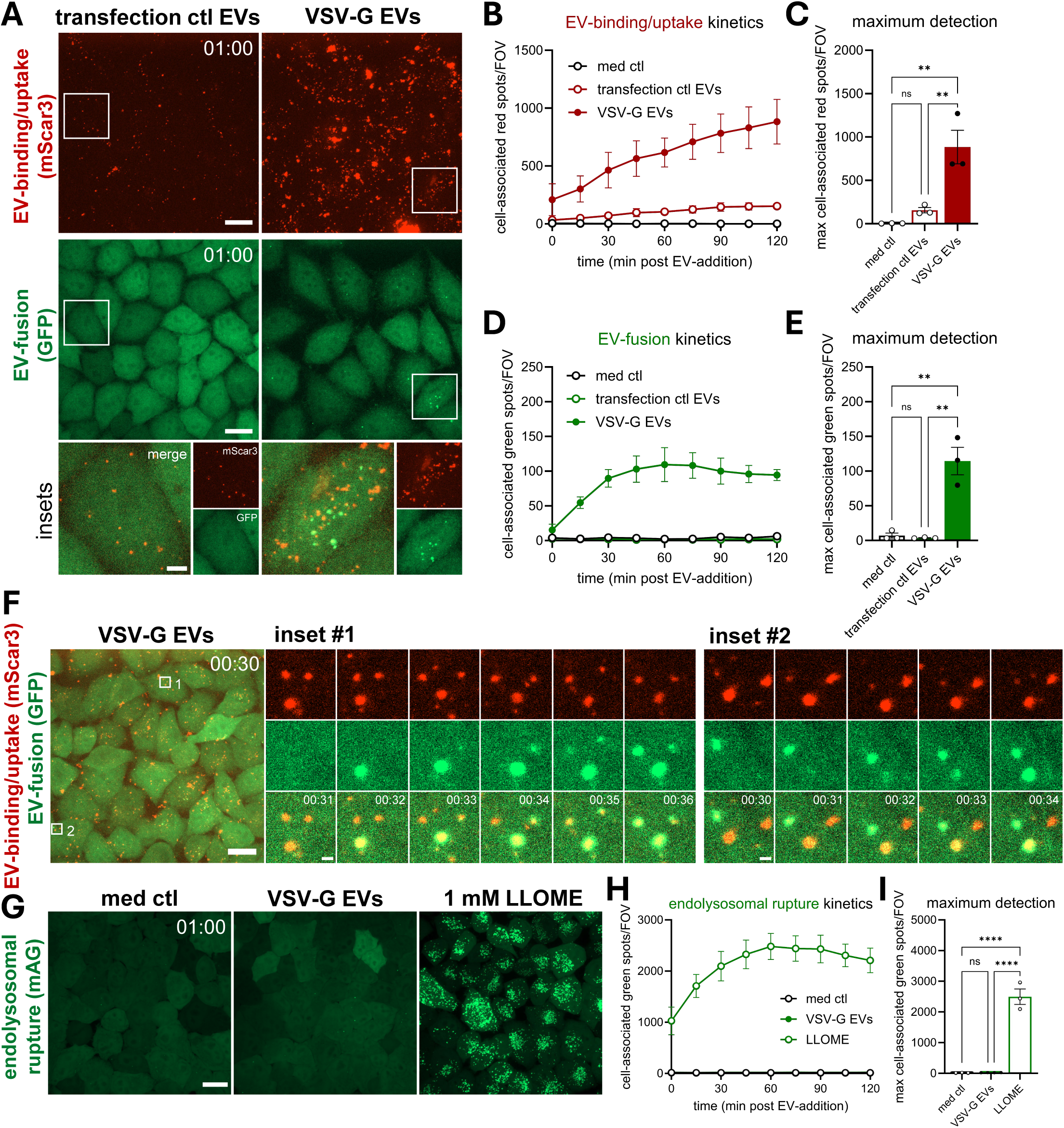
EV-FUSIM facilitates real-time visualization of VSV-G-mediated EV-fusion. **(A)** Live HeLa STAb-GFP cells were subjected to timelapse imaging immediately after addition of medium only or medium containing 10^8^ transfection ctl or VSV-G EVs, taking Z-stacks at a 15 min time interval. Images are maximum intensity projections acquired at 1h post-EV-addition, showing fluorescence channels for mScar3 **(top)** and GFP **(middle)** in the same fields-of-view. Gamma was adjusted to 2 (mScar3) or 1.2 (GFP) for visualization purposes only. Scale bars represent 20 µm. White insets correspond to magnifications **(bottom)**, which show mScar3, GFP and merged channels. Scale bar represents 5 µm. Images are representative of n=3 independent experiments. **(B-E)** After segmentation of cells in 3D based on the GFP channel, cell-associated fluorescent spots in the red and green channel were counted for each field-of-view. In tandem, the total cell volume per field-of-view was counted, which was used to correct spot counts for differences to the average cell volume per field-of-view. Graphs show the corrected mean spots per timepoint **(B, D)** or mean maximum spot detection over the course of the experiment **(C, E)** per field-of-view for both mScar3 and GFP channels ± SEM, calculated from n=3 independent experiments with 2-6 fields-of-view per condition each. * p≤0.05, ** p≤0.01 and *** p≤0.001 as determined by one-way ANOVA with Tukey’s multiple comparisons test. **(F)** Live HeLa STAb-GFP were subjected to timelapse imaging 30 min after addition of medium containing 10^8^ VSV-G EVs, taking Z-stacks at a 1 min time interval. Images are representative of n=2 independent experiments. Overview image is a maximum intensity projection of the start of the experiment, showing a merge of fluorescence channels for mScar3 and GFP. Scale bar represents 20 µm. Insets shown in white correspond to magnified images of single EV-containing endosomes on the right, showing fluorescence channels for mScar3, GFP and a merge of both at the indicated timepoints post-EV-addition. Scale bars represent 1 µm. **(G)** Live HeLa mAG-Gal3 cells were subjected to timelapse imaging immediately after addition of medium only, medium containing 10^8^ VSV-G EVs or medium containing 1 mM LLOME, taking Z-stacks at a 15 min time interval. Images are maximum intensity projections acquired at 1h post-EV-addition, showing the fluorescence channel for mAG. Gamma was adjusted to 1.2 (mAG) for visualization purposes only. Scale bar represents 20 µm. **(H, I)** After segmentation of cells in 3D based on the mAG channel, cell-associated fluorescent spots in the green channel were counted for each field-of-view. In tandem, the total cell volume per field-of-view was counted, which was used to correct spot counts for differences to the average cell volume per field-of-view. Graph shows the corrected mean spots per timepoint **(H)** or mean maximum spot detection over the course of the experiment **(I)** per field-of-view for the mAG channel ± SEM, calculated from n=3 independent experiments with 5-6 fields-of-view per condition each. **** p≤0.0001 as determined by one-way ANOVA with Tukey’s multiple comparisons test.

### EV-FUSIM can disentangle mechanisms controlling EV-binding/uptake or EV-fusion

Based on the capacity of EV-FUSIM to simultaneously detect EV-binding, -uptake and - fusion, we tested whether this technology can be used to disentangle factors that specifically control discrete steps in this cascade. As a proof-of-concept, we treated HeLa STAb recipient cells with V-ATPase inhibitor Bafilomycin A1 (BafA1). BafA1 is known to abrogate the pH-dependent fusogenic capacity of VSV-G, both in the context of infection with bona fide VSV (Burkard et al., 2014) and for VSV-G-containing EVs (Somiya & Kuroda, 2021b), by preventing endosomal acidification. Treatment with BafA1 prior to addition of EVs affected neither binding nnor uptake of VSV-G EVs, did not affect the binding/uptake of VSV-G EVs, as indicated by the lack of a significant difference in red spot detection compared to vehicle control-treated cells (**Fig. 7A-C, Suppl. Fig. 8**). In contrast, BafA1 completely abrogated the detection of fusion events, as evidenced by a reduction of green spot detection to levels observed for cells which were not incubated with EVs (**Fig. 7A, D, E**). The independent analysis of perturbations in EV-binding, - uptake or -fusion opens up unique opportunities to delineate mechanisms underlying each of these processes separately as well as explore their interconnectedness.

**Figure 7.**
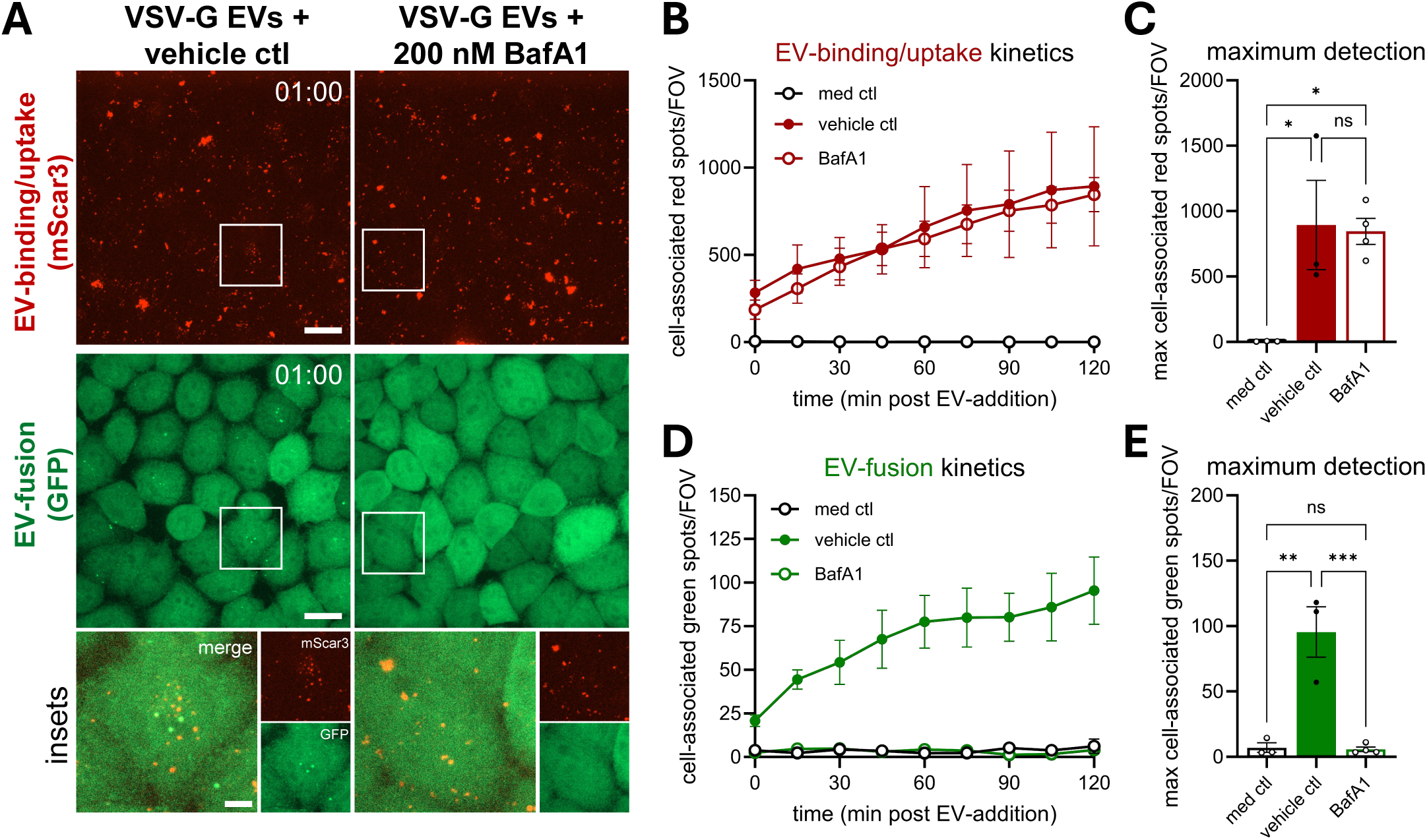
Inhibition of endosomal acidification specifically abrogates VSV-G-mediated EV-fusion without affecting EV-binding/uptake. **(A)** Live HeLa STAb-GFP cells were pre-treated with DMSO (vehicle ctl) or Bafilomycin A1 for 30 min, followed by addition of medium containing 10^8^ VSV-G EVs in the continued presence of the compounds, then immediately subjected to timelapse imaging, taking Z-stacks at a 15 min time interval. Images are maximum intensity projections acquired at 1h post-EV-addition, showing fluorescence channels for mScar3 **(top)** and GFP **(middle)** in the same fields-of-view. Gamma was adjusted to 2 (mScar3) or 1.2 (GFP) for visualization purposes only. Scale bars represent 20 µm. White insets correspond to magnifications **(bottom)**, which show mScar3, GFP and merged channels. Scale bar represents 5 µm. Images are representative of n=3-4 independent experiments. **(C-E)** After segmentation of cells in 3D based on the GFP channel, cell-associated fluorescent spots in the red and green channel were counted for each field-of-view. In tandem, the total cell volume per field-of-view was counted, which was used to correct spot counts for differences to the average cell volume per field-of-view. Graphs show the corrected mean spots per timepoint **(B, D)** or mean maximum spot detection over the course of the experiment **(C, E)** per field-of-view for both mScar3 and GFP channels ± SEM, calculated from n=3-4 independent experiments with 2-6 fields-of-view per condition each. * p≤0.05, ** p≤0.01 and *** p≤0.001 as determined by one-way ANOVA with Tukey’s multiple comparisons test.

### Subcellular localization of EV-fusion events by colocalizing with compartments of interest

Precise subcellular locations of EV-fusion remain poorly understood. We reasoned that the spatial resolution offered by EV-FUSIM could aid in understanding at which types of membranes VSV-G EVs fuse. Prior to the addition of EVs, HeLa STAb recipient cells were labelled with either CellMask, to stain the plasma membrane and endosomes originating from it, or LysoTracker, to stain acidic late endosomes and lysosomes (**Fig. 8A**) (Jenkins et al., 2015; Barral et al., 2022). These staining procedures did not negatively affect EV-fusion, indicated by similar numbers and kinetics of EV-fusion events in dye-stained and unstained control cells (**Suppl. Fig. 9**). Next, the occurrence of EV-fusion events at the stained subcellular compartments was assessed, based on fluorescent intensity measurements in the segmented 3D STAb-GFP objects (see Materials & Methods for detailed description). EV-fusion events occurred at the plasma membrane and/or endosomes, as the vast majority of spots were detected at CellMask-stained compartments (**Fig. 8B**). In contrast, EV-fusion events rarely occurred at acidic late endosomes or lysosomes across the course of the experiment, as evidenced by infrequent spot detection at LysoTracker-stained compartments (**Fig. 8C**). Indeed, the fraction of EV-fusion events occurring at the plasma membrane and/or endosomes was significantly higher than that occurring at acidic compartments (**Fig. 8D**). By differentially segmenting CellMask-positive EV-fusion events based on their proximity to the cell surface, we could further confirm that the vast majority of EV-fusion events occurred at internalized endosomal membranes, not at the plasma membrane (**Fig. 8E**). Together, these data indicate that VSV-G EVs preferentially fuse at early, less acidic endosomal compartments.

**Figure 8.**
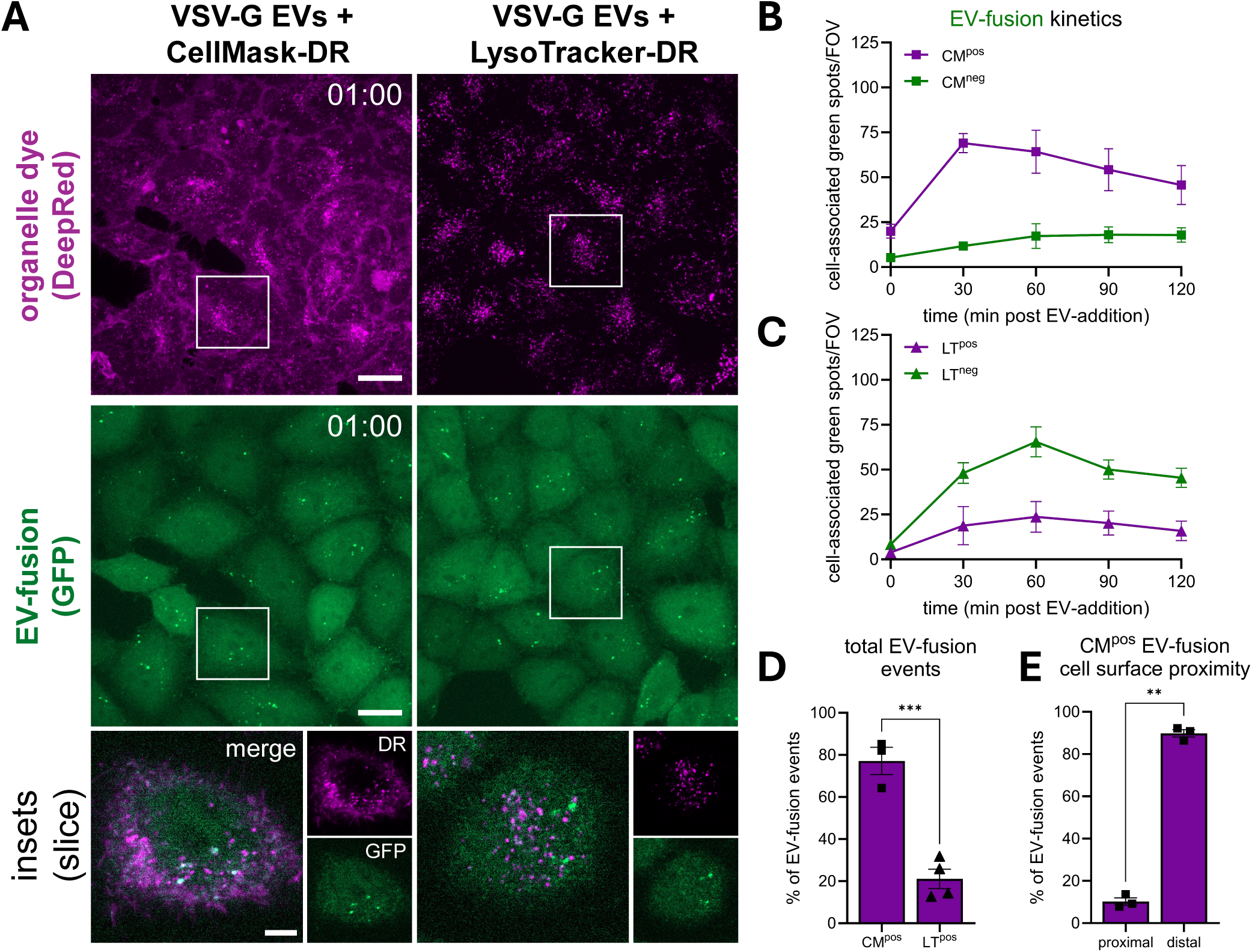
Localization analysis suggests VSV-G-mediated EV-fusion occurs predominantly in early endosomes. **(A)** Live HeLa STAb-GFP cells were left untreated or stained for 15 min with either 20 nM LysoTracker-DeepRed or 5 µg/mL CellMask, washed with medium followed by addition of 10^8^ VSV-G EVs, then immediately subjected to timelapse imaging, taking Z-stacks at a 30 min time interval. Overview images are maximum intensity projections acquired at 1h post-EV-addition, showing fluorescence channels for DeepRed **(top)** and GFP **(middle)** in the same fields-of-view. Gamma was adjusted to 1.2 (DeepRed and GFP) for visualization purposes only. Scale bars represent 20 µm. White insets correspond to single Z-slice magnifications **(bottom)**, which show DeepRed, GFP and merged channels. Scale bar represents 5 µm. Images are representative of n=3-4 independent experiments. **(B, C)** After segmentation of cells in 3D based on the GFP channel, cell-associated fluorescent spots in the green channel were counted for each field-of-view and differentially classified as dye-negative or –positive, based on a threshold of at least 25% of the mean STAb spot volume staining positive. In tandem, the total cell volume per field-of-view was counted, which was used to correct spot counts for differences to the average cell volume per field-of-view. Graphs show the corrected mean spots per timepoint, per class ± SEM, calculated from n=3-4 independent experiments with 4-6 fields-of-view per condition each. **(D)** Graph shows the mean percentage of the total GFP spots staining positive for each dye ± SEM, based on the cumulative spots over 2h per field-of-view and calculated from n=3-4 independent experiments with 4-6 fields-of-view per condition each. *** p≤0.001 as determined by unpaired t-test. **(E)** CellMask-positive GFP spots were further sub-classified based on their distance to the segmented cell surface, as proximal (<500 nm) or distal (>500 nm). Graph shows the mean percentage of the total CellMask-positive GFP spots in either class ± SEM, based on the. data shown before ** p≤0.01 as determined by paired t-test.

The subcellular location where EVs deliver their cargo may influence the likelihood that bioactive molecules contained in EVs reach their site of action. The nucleus has been suggested as an important site of action for cargo delivered by both EVs and viruses (Corbeil et al., 2020; Santos et al., 2021, 2023). EV-FUSIM offers the opportunity to assess whether the actual fusion process takes place in the proximity of specific target organelles. We used our localization analysis pipeline to differentially classify spots overlapping with a blue fluorescent protein tagged to a nuclear localization signal (NLS-BFP), to assess the proximity of EV-fusion events to the nucleus. Interestingly, a large fraction of EV-fusion events overlapped with NLS-BFP, indicating close proximity to the nucleus (**Fig. 9A**). Nucleus-proximal EV-fusion was a stable constituent of the total EV-fusion process over time, representing approximately 33.7% of the total detected events (**Fig. 9B, C**). This indicates VSV-G EVs can fuse with endosomal membranes in very close proximity to the nuclear envelope, potentially delivering cargo in a favorable location for transport into the nucleus. Combined, our data illustrate that the high spatio-temporal resolution of EV-FUSIM offers ample opportunities to unravel previously unexplored aspects of the EV-fusion process, including subcellular localization.

**Figure 9.**
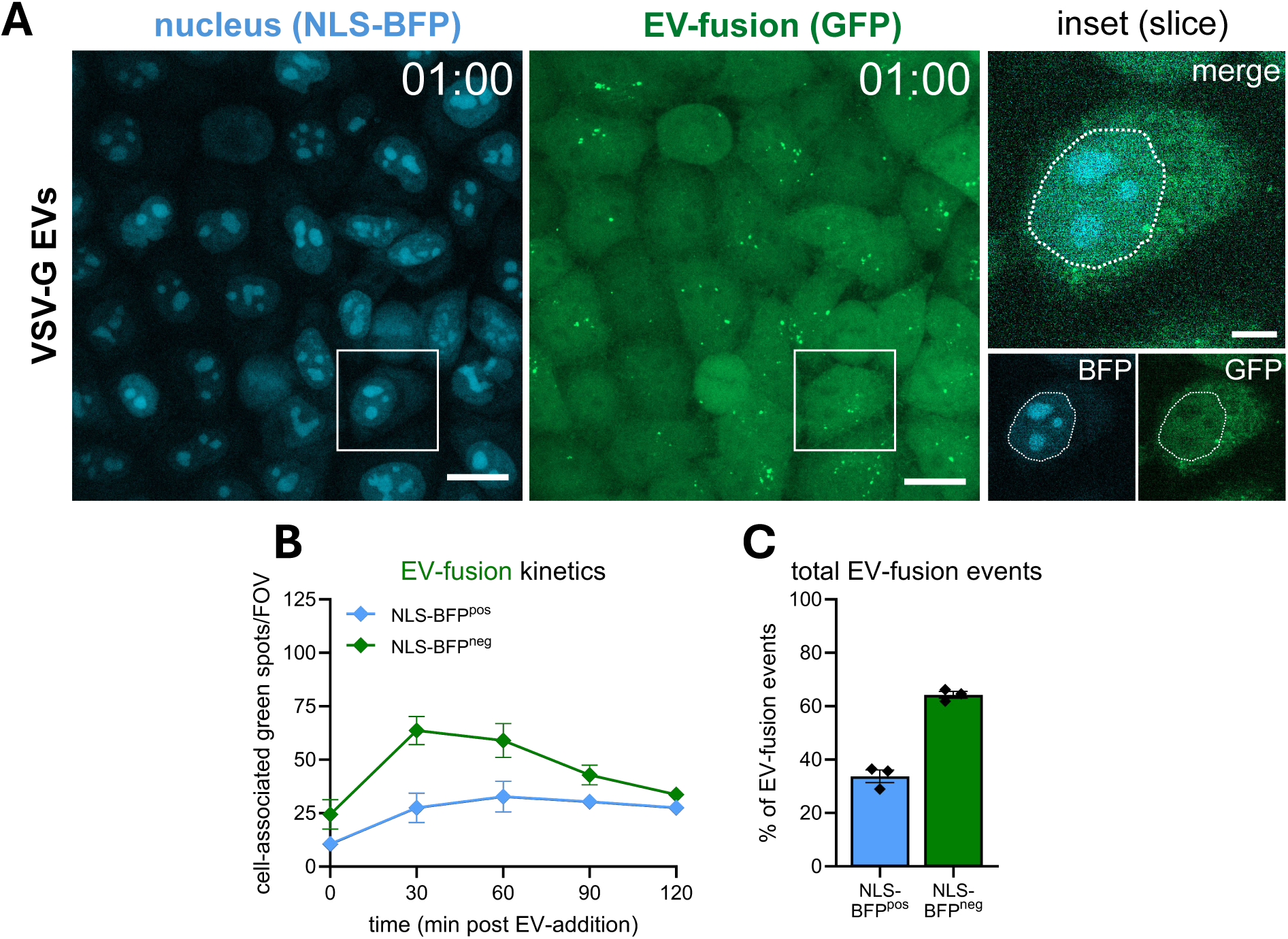
A substantial fraction of VSV-G-mediated EV-fusion events occurs in close proximity to the nucleus. **(A)** Live HeLa STAb-GFP cells expressing a nuclear localization signal-tagged BFP were subjected to timelapse imaging immediately after addition of medium containing 10^8^ VSV-G EVs, taking Z-stacks at a 30 min time interval. Overview images are maximum intensity projections acquired at 1h post-EV-addition, showing fluorescence channels for BFP **(left)** and GFP **(right)** in the same fields-of-view. Gamma was adjusted to 1.2 (BFP and GFP) for visualization purposes only. Scale bars represent 20 µm. White insets correspond to single Z-slice magnifications **(right)**, which show BFP, GFP and merged channels. Scale bar represents 5 µm. White dashed line indicates nuclear outline, drawn for visualization purposes only. Images are representative of n=3 independent experiments. **(B)** After segmentation of cells in 3D based on the GFP channel, cell-associated fluorescent spots in the green channel were counted for each field-of-view and differentially classified as BFP-negative or –positive, based on a threshold of at least 25% of the mean STAb spot volume staining positive. In tandem, the total cell volume per field-of-view was counted, which was used to correct spot counts for differences to the average cell volume per field-of-view. Graph shows the corrected mean spots per timepoint, per class ± SEM, calculated from n=3 independent experiments with 6 fields-of-view each. **(C)** Graph shows the mean percentage of the total GFP spots staining positive for BFP ± SEM, based on the cumulative spots over 2h per field-of-view and calculated from n=3 independent experiments with 6 fields-of-view each.

## Discussion

Detailed mechanistic understanding of EV-target cell interactions is a pre-requisite for exploring the role of EVs in (patho)physiological processes, modulating their functional properties, and ultimately harnessing them for therapeutic approaches. Here, we extended the EV-toolbox by developing an <u>E</u>xtracellular <u>V</u>esicle <u>Fu</u>sion <u>S</u>patiotemporal <u>Imaging</u> M<u>ethod</u> (EV-FUSIM) – a methodology for live-cell imaging of EV-binding, -uptake and -fusion with target cells in time and space.

We engineered EV-donor cells to tag the luminal EV-membrane with red fluorescent proteins coupled to SunTag peptides using a palmitoylation motif (**Fig. 2**). Additional VSV-G expression in donor cells induced the release of SunTagged, VSV-G-containing EVs (**Fig. 3**). Upon addition of SunTagged EVs to recipient cells expressing anti-SunTag single-chain antibodies, we were able to detect EV-binding and -uptake across different EV-concentrations and conditions in real-time for prolonged periods (**Fig. 4, 5**). For VSV-G EVs, we were additionally able to detect EV-fusion in live cells, (**Fig. 4, 5**). By zooming in on individual EV-containing endosomes and imaging at short time intervals, we provide real-time visualization of EV-fusion from endosomal compartments (**Fig. 6**). Using chemical inhibition of endosomal acidification, we demonstrate that simultaneous detection of EV-binding, -uptake and -fusion in real-time facilitates the disentanglement of distinct steps in the EV-target cell interaction cascade (**Fig. 7**). Lastly, localization analysis can be combined with EV-FUSIM to determine where in the endosomal system EV-fusion occurs, and which cellular organelles are in close proximity to the EV-fusion event could thus be targets of the delivered EV-cargo (**Fig. 8, 9**).

Using EV-FUSIM, we were able to comprehensively characterize the interactions of VSV-G EVs with target cells in time and space. Firstly, we demonstrate that VSV-G increases the binding and uptake efficiency of EVs (**Fig. 4-6**). Similar observations have been made previously using bioluminescent reporters or flow cytometry (Bui et al., 2023; Stranford et al., 2023) and could be linked to binding of the viral glycoprotein to its widely expressed cellular receptor, LDLR, thereby inducing its internalization (Finkelshtein et al., 2013). Secondly, we show that VSV-G-mediated EV-fusion occurs rapidly, with STAb-GFP spots visible immediately after the start of imaging and maxing out between 30 to 60 minutes after addition of EVs (**Fig. 4-6**). These rapid fusion kinetics are in line with previous reports demonstrating this for both VSV-G-containing EVs and bona fide VSV, using bioluminescent and fluorescence resonance energy transfer (FRET) reporters, respectively. (Somiya & Kuroda, 2021a; Cabot et al., 2022). The fusion detection plateau we observe after 60 minutes could reflect a technical limitation of the system. It is possible that at these later timepoints the VSV-G-containing EVs still internalize and fuse, but can no longer be detected due to saturation of the limited amount of STAb-GFP molecules in recipient cells. Alternatively, the pool of EVs containing VSV-G may internalize and fuse rapidly at early timepoints, whereas SunTagged EVs without VSV-G on their surface may internalize with slower kinetics but do not have the capacity to fuse. Thirdly, we provide novel evidence that the mechanism of VSV-G-mediated EV-cargo delivery is via EV-fusion, rather than endolysosomal rupture (**Fig. 6**). Furthermore, we demonstrate that fusion of VSV-G EVs occurs predominantly in early, less acidic endosomes, rather than in late endosomes or lysosomes (**Fig. 8**). This is in line with previous studies showing fusion-promoting conformational changes in VSV-G occur at slightly acidic pH values (around 6.2) using *in vitro* liposome fusion assays or FRET-based pH sensors (Cabot et al., 2022; Kim et al., 2017). Such pH values occur early in the endosomal system (Wang et al., 2017). Lastly, we observed that a substantial fraction of VSV-G-mediated EV-fusion events occurs in close proximity to the nucleus (**Fig. 9**). Migration of the viral G-protein to the nucleus during VSV infection was reported nearly three decades ago, although the implications for virus infection remained underexplored (Da Poian et al., 1996). Our data suggest that VSV-G EVs could be transported through the endosomal system to a similar location, and eventually fuse near the nuclear envelope. Many therapeutically relevant EV-cargoes, such as Cas9-gRNA complexes, need to reach the nucleus for their activity. Nucleus-proximal EV-fusion might be beneficial or even required for functional delivery of such cargoes, by minimizing their presence in the cytosol, where there is a potential risk of degradation and/or inactivation. Altogether, exploration of the fundamental mechanisms of VSV-G-mediated EV-cargo delivery as described here could prove invaluable in the development of novel and optimization of current VSV-G EV-based therapeutics. Real-time detection of distinct steps in the delivery pathways, as facilitated by EV-FUSIM, can in the future be used to assess the effects of e.g. VSV-G mutants or drugs, in order to ultimately improve functional cargo delivery. In addition, it may allow for successful reiteration of gain-of-function screens exploring alternative EV-fusion factors, which have thus far been largely inconclusive (Liang et al., 2025).

The live-cell imaging data we present here substantiate previous findings that the efficiency of EV-mediated cargo delivery is generally low (De Jong et al., 2020; Ilahibaks et al., 2023; Liang et al., 2025; Obuchi et al., 2025). EV-FUSIM recipient cells never showed a significant increase in green fusion spot detection when exposed to EVs lacking VSV-G, even when treated with 12,500 EVs/cell and followed for 12 hours (**Fig. 4, 5**). EVs released via different biogenesis pathways or by diverse cell types exhibit strong differences in molecular cargo, and may therefore also differ in fusogenic capacity. In this study, we used a palmitoylation motif to tag EVs through general membrane labelling, which likely captures a wide spectrum of EVs released via different biogenesis pathways. Indeed, mScarlet3 was detected on both the plasma membrane and internal compartments upon expression of the construct in HeLa cells (**Fig. 2**). Our data indicate that EVs released by HeLa cells do not fuse with HeLa target cells under the specified conditions at an efficiency high enough to detect in our assay. Nevertheless, we cannot rule out that these cells release specific subsets of highly fusogenic EVs which are either not tagged using our strategy or exhibit a fusion frequency that is lower than the throughput of our assay. In the future, mScar3-SunTag tagging of marker proteins present in specific EV-subsets combined with strategies to enrich for tagged subsets could address such limitations (Bobbili et al., 2024; Obuchi et al., 2025). The lack of data on whether and how EV-fusion is influenced by donor and recipient cell types, culture conditions, EV-tagging and -isolation methodologies, as well as reporter assays constitutes a major knowledge gap in the EV research field. Our novel technology offers the opportunity for direct real-time detection of EV-fusion with target cells to explore the influence of these variables without confounding effects of post-fusion processes.

In conclusion, an increased appreciation in the field of the tightly regulated nature of EV-fusion with target cells calls for methods that directly and specifically assess fusion efficiency with high spatiotemporal precision. The EV-FUSIM system, which we have described here, can be utilized in the future to identify fusogenic EV-subsets, fusion-promoting recipient cell types or states, and molecular players involved in these processes. Although viral fusion proteins can enhance fusion of engineered EVs, their immunogenicity is a clear disadvantage for therapeutic use. This calls for the discovery of novel fusogenic moieties or EV-fusion-enhancing drugs for use in the clinic. Being able to monitor and understand the fusion step in the EV-life cycle will be instrumental for engineering of vesicles that efficiently deliver intraluminal therapeutic payloads at the desired subcellular location.

## Author contributions

Conceptualization: J.E., K.A.Y.D., M.E.T., R.W. & E.N.M.N-‘tH. Formal analysis: J.E. & R.W. Funding Acquisition: E.N.M.N-‘tH. Investigation: J.E. Methodology: J.E., K.A.Y.D., H.H.R. & R.W. Supervision: R.W. & E.N.M.N-‘ tH. Writing – original draft: J.E. Writing – review & editing: J.E., R.W. & E.N.M.N-‘tH.

## Acknowledgements

We are grateful to the Centre for Cellular Imaging Utrecht and the Flow Cytometry and Cell Sorting Facility, both at the Faculty for Veterinary Medicine of Utrecht University, for allowing access to their facilities, training and support. We thank dr. Pieter Vader (UMC Utrecht) for providing an antibody targeting Calnexin.

---

**Suppl. Figure 1.**
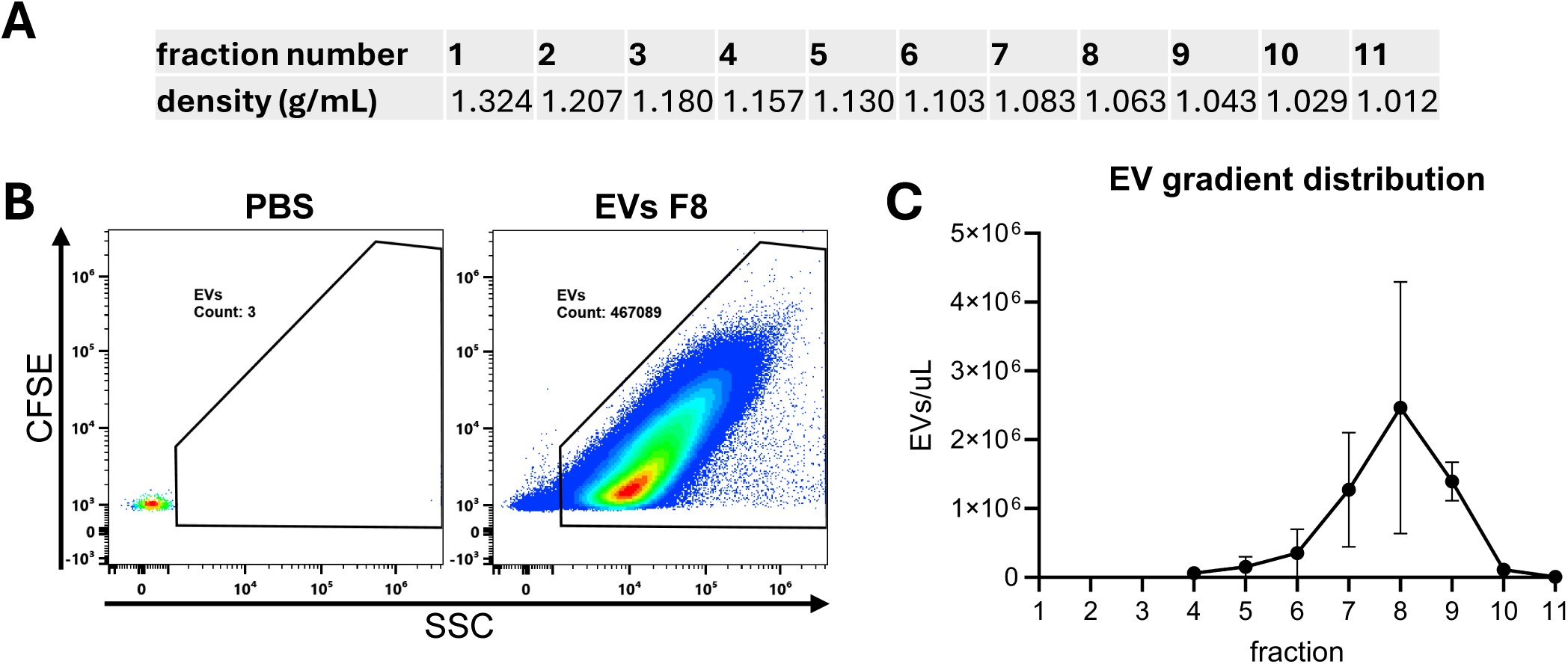
Density gradient distribution of EVs. Refractive indices of individual density gradient fractions were measured using a refractometer and converted into density values. **(B)** Dot plots showing the CFSE fluorescence vs side-scatter (SSC) profiles of PBS only and EVs in fraction 8, with representative gating strategy. **(C)** EV-concentration as measured in the gate as set in (B) across fractions 4 through 11, corrected for dilution factors to yield the EV-concentration in the fractions. Graph shows the mean EV-concentration ± SD, calculated from n=2 independent experiments.

**Suppl. Figure 2.**
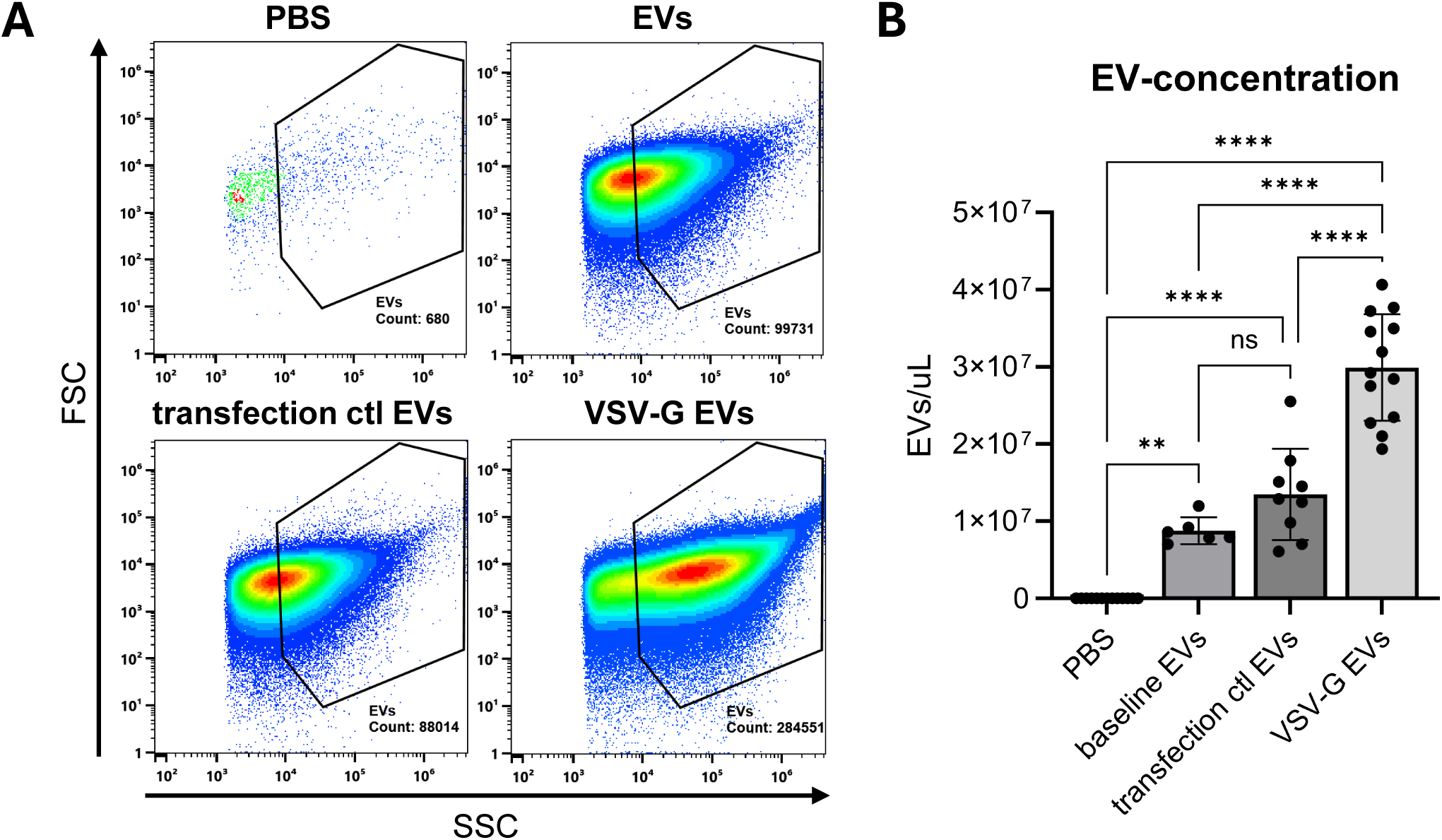
EV concentration measurements in purified EV preparations. **(A)** Dot plots showing the forward (FSC) vs side-scatter (SSC) profiles of PBS only and gradient-purified EVs isolated from the indicated cell conditions, with representative gating strategy. **(B)** EV-concentration as measured in the gate as set in (A), corrected for dilution factors to yield the EV-concentrations in the preparations. Graph shows the mean EV-concentration ± SD, calculated from n=6-12 independent experiments. ** p≤0.01 and **** p≤0.0001 as determined by one-way ANOVA with Tukey’s multiple comparisons test.

**Suppl. Figure 3.**
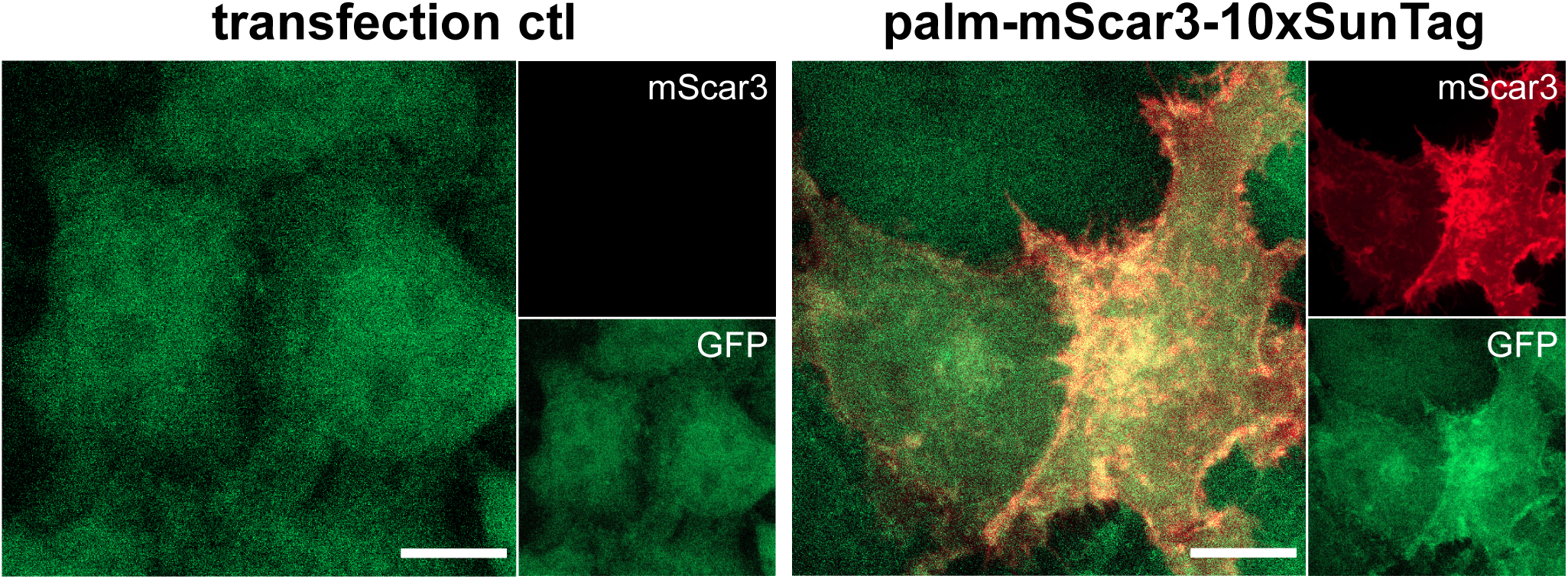
Palm-mScar3-10xST recruits STAb-GFP upon co-expression. **(A)** Live HeLa STAb-GFP cells were subjected to live-cell imaging after overnight mock- or palm-mScar3-10xST transfection. Images shown are maximum intensity projections, showing fluorescence for mScar3 (red) and GFP (green). Scale bars represent 10 µm.

**Suppl. Figure 4.**
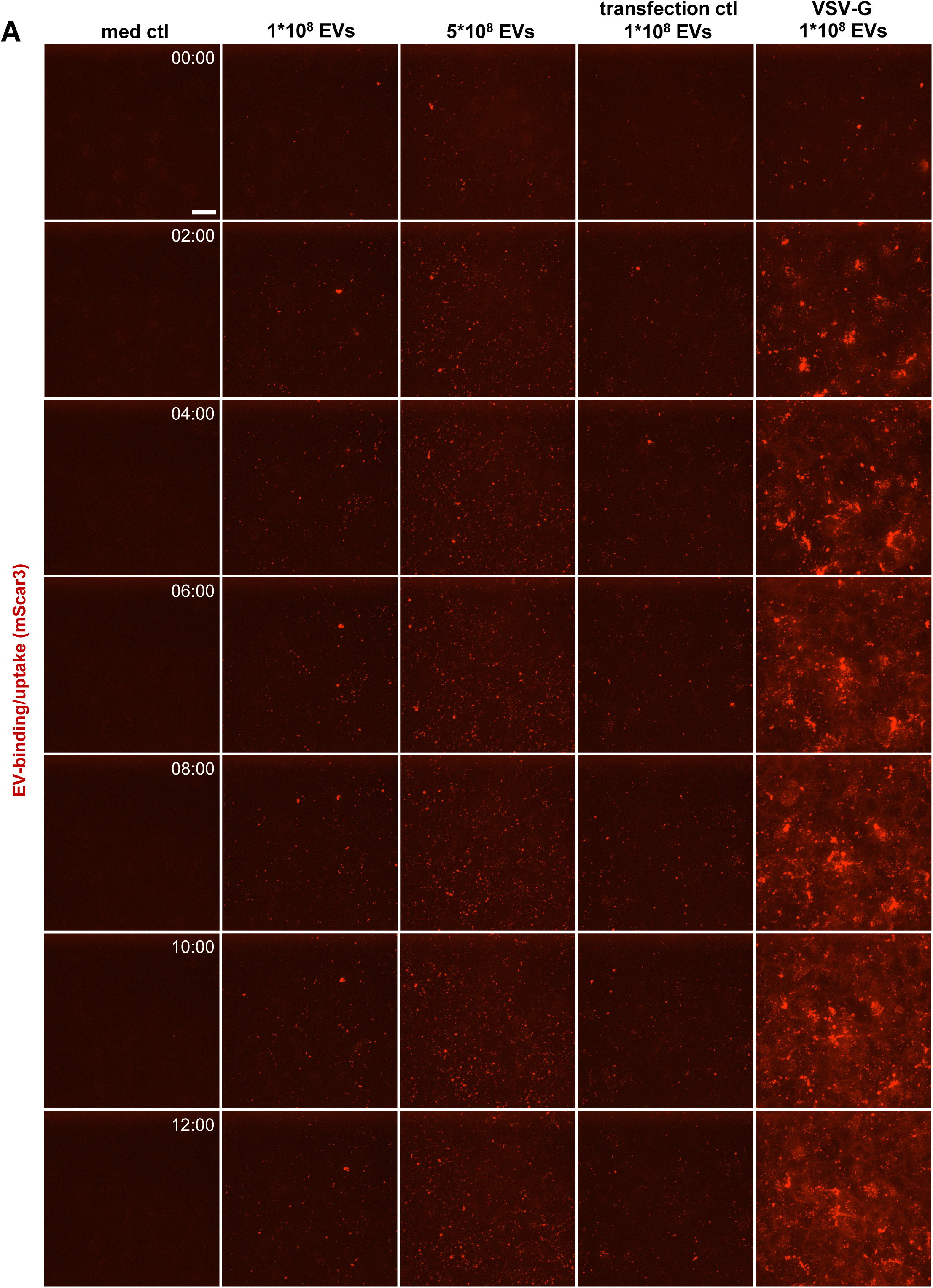
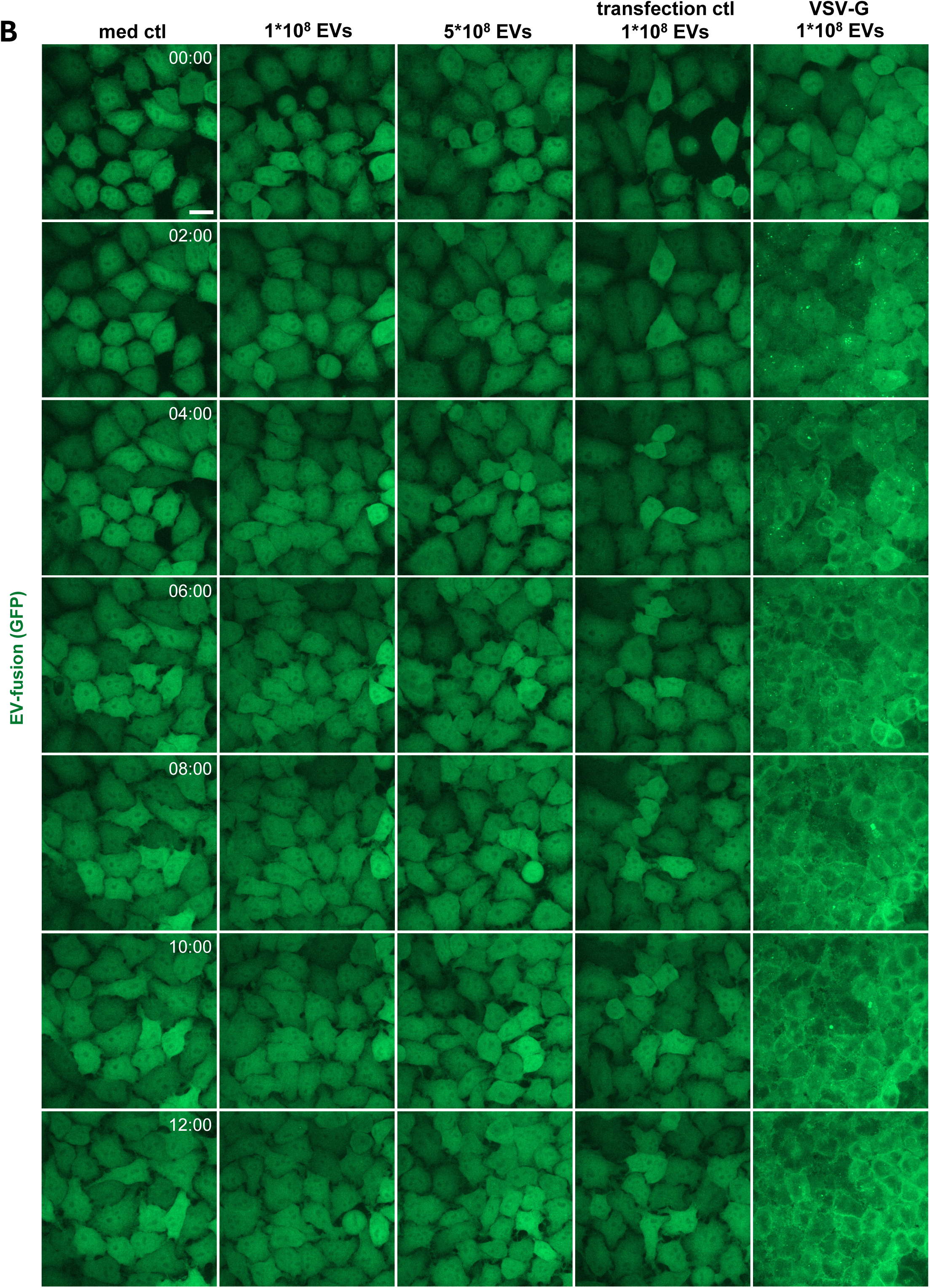
Live-cell imaging of HeLa STAb-GFP cells after addition of SunTagged EVs allows for tracing of EV-binding, -uptake and -fusion over time – extended data. **(A,B)** Additional timepoints of the data shown in Fig. 4B. Images are maximum intensity projections acquired at the indicated timepoints post-EV-addition, showing fluorescence channels for mScar3 **(A)** and GFP **(B)** in the same fields-of-view. Gamma was adjusted to 2 (mScar3) or 1.2 (GFP) for visualization purposes only. Scale bars represent 20 µm. Images are representative of n=3-4 independent experiments.

**Suppl. Figure 5.**
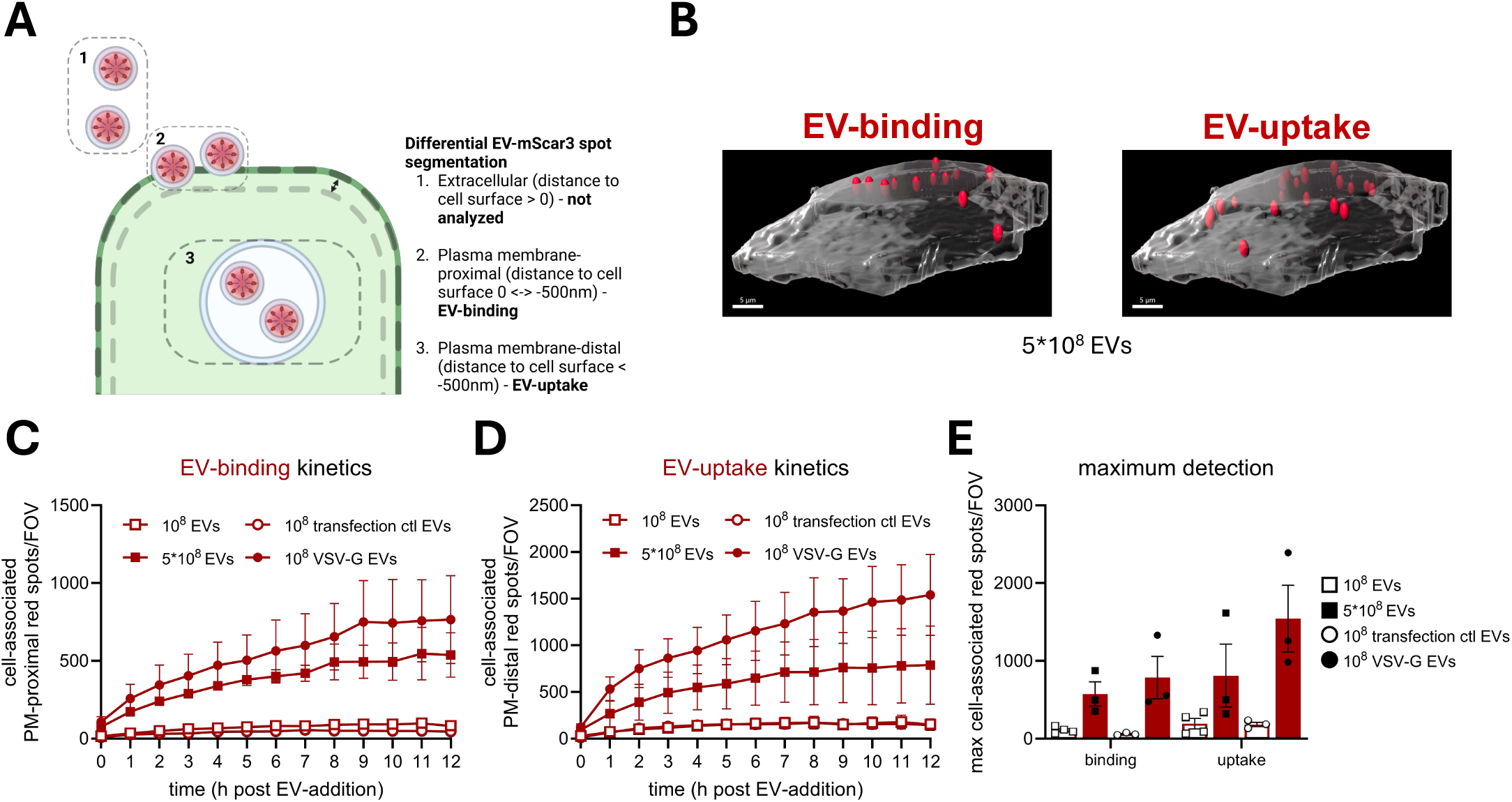
Differential segmentation allows for separate quantification of EV-binding and -uptake. **(A)** Schematic overview of differential segmentation procedure. See Materials & Methods for detailed description. Created in BioRender. van den Ende, J. (2025) https://BioRender.com/ycxyw00. **(B)** 3D rendering of cell surface as segmented in Imaris, for a single STAb-GFP cell from the experiment also shown in Fig. 4. Cell boundary is depicted in gray. Differentially segmented mScar3 spots are shown in red, with a distance of <500 nm (EV-binding, left) or >500 nm (EV-uptake, right) to the segmented cell surface. Scale bars represent 5 µm. **(C-E)** Quantification of the data shown in Fig. 5 incorporating differential segmentation for EV-binding and -uptake, corrected for differences to the average cell volume per field-of-view. Graphs show the corrected mean mScar3 spots per timepoint **(C,D)** or mean maximum spot detection over the course of the experiment **(E)** per field-of-view for EV-binding or EV-uptake ± SEM, calculated from n=3-4 independent experiments with 3-6 fields-of-view per condition each.

**Suppl. Figure 6.**
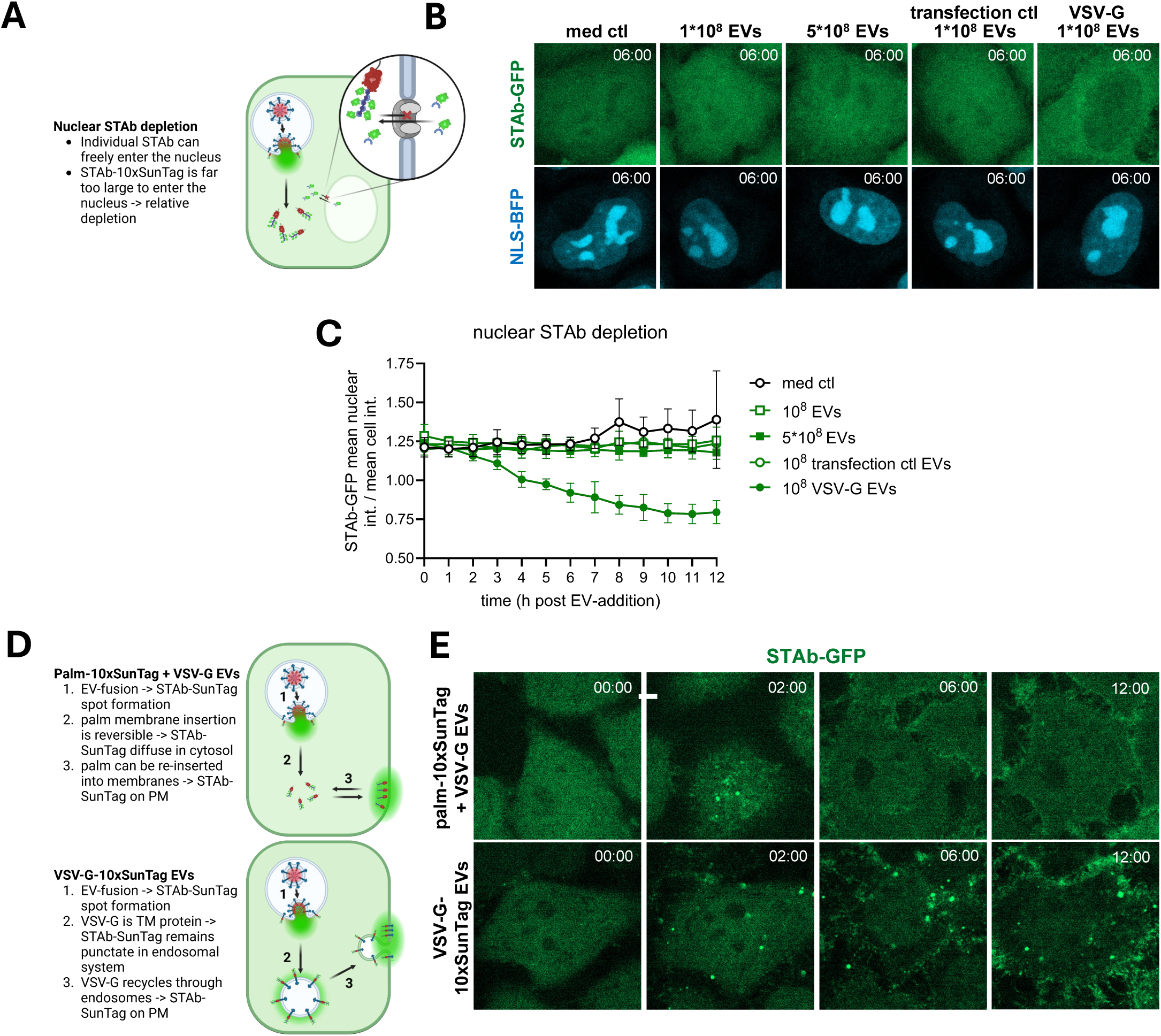
STAb-GFP localization post-EV-fusion. **(A)** Schematic depiction of observed post-fusion depletion of STAb-GFP from the nucleus. As described before, sequestration of STAb-GFP by SunTag in the cytosol results in a relative depletion of the nuclear signal, as the large complexes formed by interaction of multiple STAb molecules with the SunTagged molecule prohibits shuttling through nuclear pores. Created in BioRender. van den Ende, J. (2025) https://BioRender.com/5yc11ya. **(B)** Experimental data supporting model in (A). Live HeLa STAb-GFP cells expressing a nuclear localization signal-tagged blue fluorescent protein (NLS-BFP) were subjected to timelapse imaging immediately after addition of medium only or medium containing the indicated concentrations and types of EVs, taking Z-stacks at a 1h time interval. Images are mid-nuclear Z-slices acquired at 6h post-EV-addition, showing fluorescence channels for GFP **(top)** and BFP **(middle)** in the same fields-of-view. Scale bars represent 5 µm. **(C)** After segmentation of cells in 3D based on the GFP channel, a nuclear volume was segmented based on the BFP channel. GFP fluorescent intensity was measured in both the total cell and nuclear volumes. Graph shows the calculated ratio of the mean nuclear intensity to the mean total cell intensity ± SD, calculated from 6 fields-of-view per condition. **(D)** Schematic depiction of observed post-fusion localization of STAb-GFP, reflecting the sorting of the SunTagged EV-associated molecule. When using palm-10xST EVs which are additionally mounted with VSV-G, as described in the rest of the data, STAb-GFP puncta eventually disappear. This is likely the result of a balance between palmitoylation and depalmitoylation in the target cell, resulting in diffusion of STAb-SunTag complexes into the cytosol. Eventually, these complexes can be seen to be re-inserted at the plasma membrane. When using EVs carrying directly SunTagged VSV-G, STAb-GFP remains punctate post-fusion, indicating it recycles through the endosomal system as is known for VSV-G, eventually concentrating at the plasma membrane. Created in BioRender. van den Ende, J. (2025) https://BioRender.com/3rw27gl. **(E)** Experimental data supporting models in (D). Live HeLa STAb-GFP cells were subjected to timelapse imaging immediately after addition of medium containing EVs isolated from VSV-G-transfected HeLa palm-mScar3-10xST cells, or from VSV-G-mScar3-10xST-transfected HeLa WT cells. Images are mid-nuclear Z-slices acquired at the indicated times post-EV-addition, showing GFP fluorescence. Scale bars represent 5 µm.

**Suppl. Figure 7.**
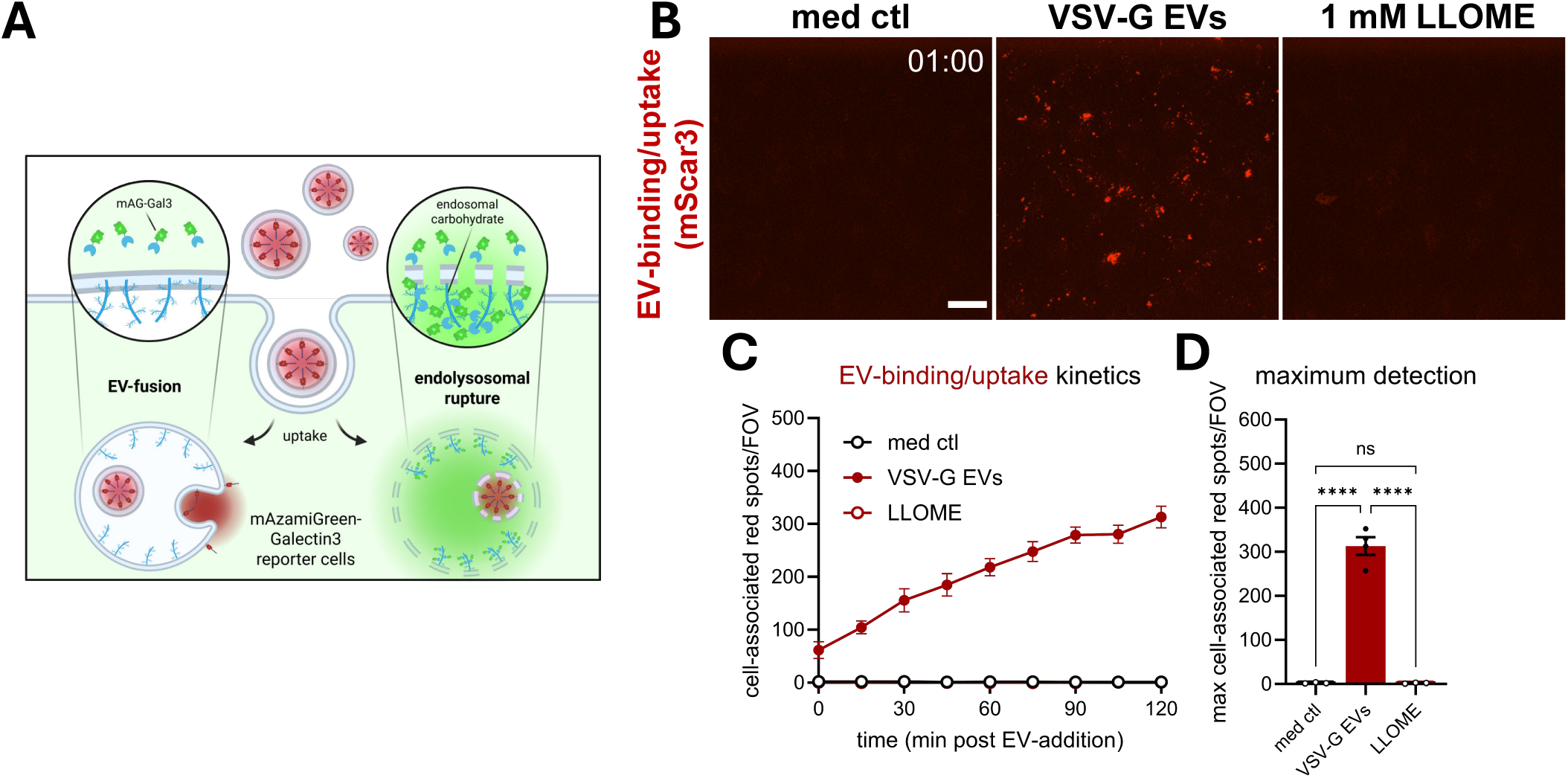
VSV-G-mediated EV-fusion occurs in the absence of endolysosomal rupture – extended data. **(A)** Schematic of Galectin-3 reporter assay principle. HeLa cells were engineered to express mAzamiGreen-tagged Galectin-3, which is present abundantly in the cytosol and has strong affinity for carbohydrate moieties. Many of these are present inside the endosomal lumen, but this pool remains inaccessible for cytosolic Gal3 unless there is rupture of the endosomal membrane. Upon addition of VSV-G EVs, red spots will indicate EV-binding/uptake, whereas green spots will indicate endolysosomal rupture. As we know SunTag is exposed to the cytosol from these EVs, absence of endolysosomal rupture detection will further prove fusion as the responsible mechanism. Created in BioRender. van den Ende, J. (2025) https://BioRender.com/tm3vcwj. **(B)** Images are maximum intensity projections acquired at 1h post-EV-addition, showing fluorescence for mScar3 in the same fields-of-view as shown in Fig. 6H. Gamma was adjusted to 2 (mScar3) for visualization purposes only. Scale bars represent 20 µm. Images are representative of n=3-4 independent experiments. **(C, D)** After segmentation of cells in 3D based on the mAG channel, cell-associated fluorescent spots in the red channel were counted for each field-of-view. In tandem, the total cell volume per field-of-view was counted, which was used to correct spot counts for differences to the average cell volume per field-of-view. Graph shows the corrected mean spots per timepoint **(H)** or mean maximum spot detection over the course of the experiment **(I)** per field-of-view for the mScar3 channel ± SEM, calculated from n=3 independent experiments with 5-6 fields-of-view per condition each. **** p≤0.0001 as determined by one-way ANOVA with Tukey’s multiple comparisons test.

**Suppl. Figure 8.**
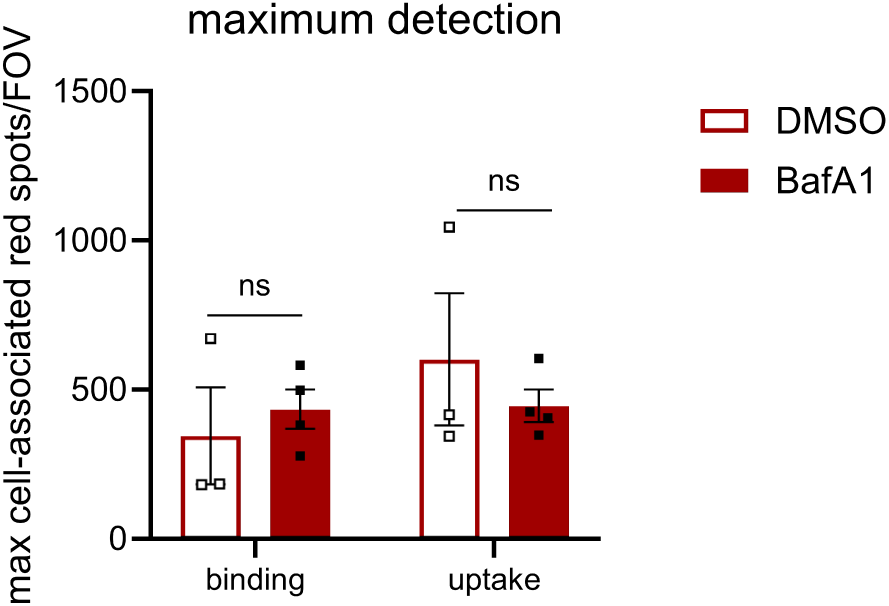
Separate quantification of EV-binding and –uptake confirms BafA1 treatment did not affect either. Quantification of the data shown in Fig. 7 incorporating differential segmentation for EV-binding and -uptake, corrected for differences to the average cell volume per field-of-view. Graphs show the corrected mean maximum mScar3 spot detection over the course of the experiment per field-of-view for EV-binding or EV-uptake ± SEM, calculated from n=3-4 independent experiments with 4-6 fields-of-view per condition each. ns p>0.05 as determined by as determined by unpaired t-test (performed separately for binding and uptake).

**Suppl. Figure 9.**
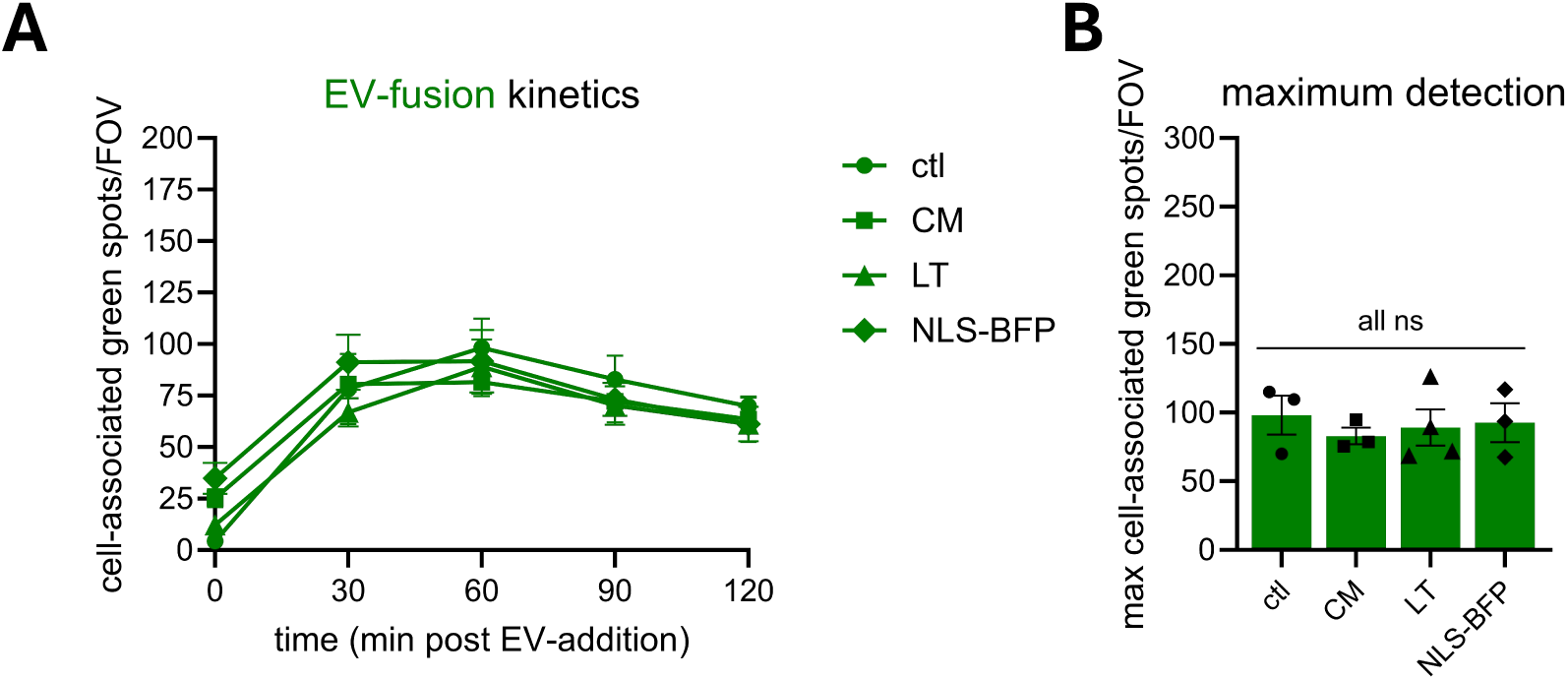
Localization experimental procedures did not alter VSV-G-mediated EV-fusion dynamics. **(A, B)** Analysis of experimental conditions also shown in Fig. 8 & 9. After segmentation of cells in 3D based on the GFP channel, cell-associated fluorescent spots in the green channel were counted for each field-of-view. In tandem, the total cell volume per field-of-view was counted, which was used to correct spot counts for differences to the average cell volume per field-of-view. Graphs show the corrected mean spots per timepoint **(A)** or mean maximum spot detection over the course of the experiment **(B)** per field-of-view for the GFP channel ± SEM, calculated from n=3-4 independent experiments with 4-6 fields-of-view per condition each. ns p>0.05 as determined by one-way ANOVA with Tukey’s multiple comparisons test.

**Suppl. Video 1. EV-FUSIM facilitates real-time visualization of VSV-G-mediated EV-fusion.** Timelapse video of data also shown in Fig. 6. Live HeLa STAb-GFP cells were subjected to timelapse imaging immediately after addition of medium containing 10^8^ VSV-G EVs, taking Z-stacks at a 15 min time interval. Images are maximum intensity projections acquired at a 15 min time interval, showing fluorescence channels for mScar3 **(left)** and GFP **(right)** in the same fields-of-view. Scale bars represent 20 µm. Time after EV-addition is indicated as hh:mm.

**Suppl. Table 1. Imaris object creation scripts.** Shown are the parameters used in Imaris to generate the denoted 3D objects, used for downstream analysis and quantification.

**Suppl. Table 2. Completed MIFlowCyt checklist.** Contains a completed checklist reporting on experimental variables related to flow cytometry experiments.

**Suppl. Table 3. Completed MIFlowCyt-EV checklist.** Contains an additional completed checklist reporting on additional experimental variables related to EV-flow cytometry experiments.

