## Supplemental Table 1 for "Development of a live-cell imaging assay to elucidate spatiotemporal dynamics of extracellular vesicle fusion with target cells"

*Imaris analysis scripts for object creation*

| **STAb cell Surface** | **Gal3 cell Surface** | **STEV Spots** | **STAb Spots** | **Gal3 Spots** | **CellMask Surface** | **LysoTracker Surface** | **NLS-BFP Surface** |
|---|---|---|---|---|---|---|---|
| [Algorithm]<br>Enable Region Of Interest = false<br>Enable Region Growing = true<br>Enable Tracking = true<br>Enable Classify = false<br>Enable Shortest Distance = true<br>[Segmentation Setup]<br>Source Channel Index = 1<br>Enable Smooth = true<br>Surface Grain Size = 0.600 µm<br>[Machine Learning Training]<br>All Channels = false<br>[Split]<br>Region Growing Estimated Diameter = 12.0 µm<br>[Filter Seed Points]<br>"Quality" above 0.0500<br>[Filter Surfaces]<br>"Area" above 330 um^2<br>"Intensity StdDev Ch=1 Img=1" below 4.00<br>"Intensity Mean Ch=1 Img=1" between 1.00 and 8.50<br>[Tracking]<br>Algorithm Name = Brownian Motion<br>MaxDistance = 10.0 µm<br>MaxGapSize = 3<br>Fill Gap Enable = false<br>[Filter Tracks] | [Algorithm]<br>Enable Region Of Interest = false<br>Enable Region Growing = true<br>Enable Tracking = true<br>Enable Classify = false<br>Enable Shortest Distance = true<br>[Segmentation Setup]<br>Source Channel Index = 1<br>Enable Smooth = true<br>Surface Grain Size = 0.700 µm<br>[Machine Learning Training]<br>All Channels = false<br>[Split]<br>Region Growing Estimated Diameter = 15.0 µm<br>[Filter Seed Points]<br>"Quality" above 0.0250<br>[Filter Surfaces]<br>"Volume" above 1700 um^3<br>"Intensity Median Ch=1 Img=1" between 2.00 and 180.0<br>[Tracking]<br>Algorithm Name = Brownian Motion<br>MaxDistance = 10.0 µm<br>MaxGapSize = 2<br>Fill Gap Enable = false<br>[Filter Tracks] | [Algorithm]<br>Enable Region Of Interest = false<br>Enable Region Growing = true<br>Enable Tracking = false<br>Enable Classify = true<br>Enable Region Growing = true<br>Enable Shortest Distance = true<br>[Source Channel]<br>Source Channel Index = 2<br>Estimated XY Diameter = 0.900 µm<br>Estimated Z Diameter = 1.80 µm<br>Background Subtraction = true<br>[Filter Spots]<br>"Quality" above 1.75<br>"Shortest Distance to Surfaces Surfaces=**Cell Surface (STAb or Gal3)"** below 0.00<br>[Spot Region Type]<br>Region Growing Type = Local Contrast<br>[Spot Regions]<br>Region Growing Automatic Treshold = false<br>Region Growing Manual Threshold = 0.1<br>Region Growing Diameter = Diameter From Border<br>Create Region Channel = false<br>[Classification]<br>Group Name = PMProxClass<br>Input = All Spot<br>No. of Classes = 2<br>Class:: Name = PMProxSTEV<br>Class:: Name = PMDistSTEV<br>FilterType = Filter1D<br>Type = Shortest Distance to Surfaces<br>Surfaces=CellSurfaceGal3 JE-Gal3batchFinal.icsx_[ibrx_2025-06-13T14-50-49.726]<br>Threshold 1 = -0.500 | [Algorithm]<br>Enable Region Of Interest = false<br>Enable Region Growing = true<br>Enable Tracking = false<br>Enable Classify = false<br>Enable Region Growing = true<br>Enable Shortest Distance = true<br>[Source Channel]<br>Source Channel Index = 1<br>Estimated XY Diameter = 0.850 µm<br>Estimated Z Diameter = 1.70 µm<br>Background Subtraction = true<br>[Filter Spots]<br>"Quality" above 3.50<br>"Intensity Max Ch=1 Img=1" above 20.0<br>"Shortest Distance to Surfaces Surfaces=**STAb cell Surface**" below 0.00 um<br>[Spot Region Type]<br>Region Growing Type = Local Contrast<br>[Spot Regions]<br>Region Growing Automatic Treshold = false<br>Region Growing Manual Threshold = 0.1<br>Region Growing Diameter = Diameter From Border<br>Create Region Channel = false | [Algorithm]<br>Enable Region Of Interest = false<br>Enable Region Growing = true<br>Enable Tracking = false<br>Enable Classify = false<br>Enable Region Growing = true<br>Enable Shortest Distance = true<br>[Source Channel]<br>Source Channel Index = 2<br>Estimated XY Diameter = 0.850 µm<br>Estimated Z Diameter = 1.70 µm<br>Background Subtraction = true<br>[Filter Spots]<br>"Quality" above 35.0<br>"Intensity Max Ch=2 Img=1" above 20.0<br>"Intensity StdDev Ch=2 Img=1" above 35.0<br>"Shortest Distance to Surfaces Surfaces=**Gal3 cell Surface**" below 0.00 um<br>[Spot Region Type]<br>Region Growing Type = Local Contrast<br>[Spot Regions]<br>Region Growing Automatic Treshold = false<br>Region Growing Manual Threshold = 0.1<br>Region Growing Diameter = Diameter From Border<br>Create Region Channel = false | preproc backgr filterwidth 0.3<br>[Algorithm]<br>Enable Region Of Interest = false<br>Enable Region Growing = false<br>Enable Tracking = false<br>Enable Classify = false<br>Enable Shortest Distance = false<br>[Segmentation Setup]<br>Source Channel Index = 2<br>Enable Smooth = true<br>Surface Grain Size = 0.130 µm<br>Enable Eliminate Background = false<br>[Threshold]<br>Active Threshold = true<br>Enable Automatic Threshold = false<br>Manual Threshold Value = 7.5<br>Active Threshold B = false<br>[Filter Surfaces]<br>"Number of Voxels Img=1" above 10.0 | preproc backgr filterwidth 0.3<br>[Algorithm]<br>Enable Region Of Interest = false<br>Enable Region Growing = false<br>Enable Tracking = false<br>Enable Classify = false<br>Enable Shortest Distance = false<br>[Segmentation Setup]<br>Source Channel Index = 2<br>Enable Smooth = true<br>Surface Grain Size = 0.130 µm<br>Enable Eliminate Background = false<br>[Threshold]<br>Active Threshold = true<br>Enable Automatic Threshold = false<br>Manual Threshold Value = 2<br>Active Threshold B = false<br>[Filter Surfaces]<br>"Number of Voxels Img=1" above 10.0 | [Algorithm]<br>Enable Region Of Interest = false<br>Enable Region Growing = true<br>Enable Tracking = false<br>Enable Classify = false<br>Enable Shortest Distance = false<br>[Segmentation Setup]<br>Source Channel Index = 2<br>Enable Smooth = true<br>Surface Grain Size = 0.250 µm<br>[Machine Learning Training]<br>All Channels = false<br>[Split]<br>Region Growing Estimated Diameter = 12.0 µm<br>[Filter Seed Points]<br>[Filter Surfaces]<br>"Number of Voxels Img=1" above 10.0 |
