## Supplemental Table 2 for "Development of a live-cell imaging assay to elucidate spatiotemporal dynamics of extracellular vesicle fusion with target cells"

| Requirement | Please Include Requested Information |
|---|---|
| 1.1. Purpose | To assess the distribution of CFSE-stained extracellular vesicles (EVs) in density gradients in relation to Western blot data, in order to corroborate EV-associated release of SunTag from HeLa palm-mScar3-10xST cells. Additionally, to assess the relative concentrations of purified EV-preparations isolated from differently treated HeLa WT and palm-mScar3-10xST cells, in order to normalize the amount of EVs added to recipient cells in live-cell imaging experiments across conditions and experiments. |
| 1.2. Keywords | Extracellular vesicles; EVs. |
| 1.3. Experiment variables | WT and genetically engineered cells; mock- or VSV-G-transfected cells. |
| 1.4. Organization name and address | Utrecht University, Yalelaan 1, Utrecht, The Netherlands. |
| 1.5. Primary contact name and email address | Jasper van den Ende, (first author)<br>Esther Nolte-'t Hoen, (senior author) |
| 1.6. Date or time period of experiment | 2024 – 2025 |
| 1.7. Conclusions | We developed a highly sensitive imaging method to detect EV-fusion events in real time, which has the power to illuminate the underexplored spatiotemporal dynamics of EV-fusion. |
| 1.8. Quality control measures | Buffer noise reduction; reduction of unbound fluorescent dye; optimization of sample dilution; equal volume analysis for relative particle concentration comparison. |
| 2.1.1.1. (2.1.2.1., 2.1.3.1.) Sample description | Extracellular vesicles isolated from cell culture supernatants, present in individual Optiprep density gradient fractions or as a purified preparation derived from pooled density gradient fractions. |
| 2.1.1.2. Biological sample source description | HeLa R19 (human cervical carcinoma, ATCC CCL-2) and derivative transgenic HeLa palm-mScar3-10xST cell lines. |
| 2.1.1.3. Biological sample source organism description | *Homo sapiens* (human), cervical adenocarcinoma. |
| 2.1.2.2. Environmental sample location | N.A. |
| 2.3. Sample treatment description | EVs were isolated from untreated or pCMV-VSVG-mScar3-10xST-transfected HeLa R19 cells, or from untreated or mock-transfected or pCMV-VSVG-transfected HeLa palm-mScar3-10xST cells. |
| 2.4. Fluorescence reagent(s) description | CFSE (carboxyfluorescein succinimidylester) |
| 3.1. Instrument manufacturer | Cytek |
| 3.2. Instrument model | Aurora |
| 3.3. Instrument configuration and settings | 3L 16V-14B-8R, with Enhanced Small Particle (ESP) detection configuration |
| 4.1. List-mode data files | N.A. |
| 4.2. Compensation description | N.A. |
| 4.3. Data transformation details | Events detected inside the EV-gate for PBS were subtracted from the events detected for other samples, to compensate for the buffer background. |
| 4.4.1. Gate description | Representative images of gating strategy, as applied to all conditions, are shown in the associated publication. Gating was either on SSC/CFSE axes, or on SSC/FSC axes, depending on the experiment. |
| 4.4.2. Gate statistics | See publication. |
| 4.4.3. Gate boundaries | See publication. |

**Notes**

Feel free to use more space than allocated.

You can embed graphics/figures in this document, if needed.
