## Supplemental Table 3 for "Development of a live-cell imaging assay to elucidate spatiotemporal dynamics of extracellular vesicle fusion with target cells"

| Framework Criteria | What to report | Please complete each criterion |
|---|---|---|
| 1.1 Preanalytical variables conforming to MISEV guidelines. | Preanalytical variables relating to EV sample including source, collection, isolation, storage, and any others relevant and available in the performed study. | Confluent cell layers were washed three times with PBS +Ca +Mg and incubated with cell culture medium containing 10% EV-depleted FBS. To generate this EV-depleted FBS, FBS was prediluted 1:3 in DMEM, ultracentrifuged for 16-20h at 28,000 rpm in an SW32 rotor (k-factor 256.8) and passed through a 0.22 µm filter. 24h after replacing the medium, cell culture supernatants were harvested and centrifuged for 10 min at 200xg and 2x 10 min at 500g to remove cells and cell debris. Cleared supernatant samples were treated with 5 mM MgCl2 and 0.1 mg/mL DNase I (Roche) for 30 min at 37°C, after which they were supplemented with 25 mM HEPES (Sigma-Aldrich). EVs were enriched from clarified supernatants by UC pelleting for 65 min at 28,000 rpm in an SW32 rotor (k-factor 256.8) and resuspended PBS +0.1% BSA (cleared from aggregates by ultracentrifugation for 16-20h at 100,000xg). Next, EVs were mixed with iodixanol (Optiprep; Axis-Shield, Oslo, Norway) to a final concentration of 45% and overlaid with a linear gradient of 40-5% iodixanol in PBS by adding 8 layers of 1.3 mL, decreasing in density at 5% decrements. Density gradients were centrifuged at 192,000xg for 16h in a SW41 rotor (k-factor 124). Gradient fractions of 1 mL were collected from the top and their densities assessed by refractometry. For experiments with purified EVs, gradient fractions 6-9 were diluted in 30 mL PBS +0.1% BSA (cleared from aggregates), isolated by UC pelleting for 90 min at 28,000 rpm in an SW32 rotor (k-factor 256.8) and resuspended in PBS. All samples for flow cytometry were measured fresh, immediately after finishing sample preparation/isolation. |
| 1.2 Experimental design according to MIFlowCyt guidelines. | EV-FC manuscripts should provide a brief description of the experimental aim, keywords, and variables for the performed FC experiment(s) using MIFlowCyt checklist criteria: 1.1, 1.2, and 1.3, respectively. Template found at [www.evflowcytometry.org.](www.evflowcytometry.org) | 1.1 Aim) To assess the distribution of CFSE-stained EVs in density gradients in relation to Western blot data, in order to corroborate EV-associated release of SunTag from HeLa palm-mScar3-10xST cells. Additionally, to assess the relative concentrations of purified EV-preparations isolated from differently treated HeLa WT and palm-mScar3-10xST cells, in order to normalize the amount of EVs added to recipient cells in live-cell imaging experiments across conditions and experiments. **1.2 Keywords)** EVs; extracellular vesicles, SunTag;ST. **1.3 Experimental variables)** EVs were isolated from untreated, mock-transfected and VSV-G-transfected HeLa WT and palm-mScar3-10xST cells. Fluorescence-based triggering was used for particle thresholding in gradient fractions, whereas scatter-based triggering was used in purified EV-preparations. |
| 2.1 Sample staining details | State any steps relating to the staining of samples. Along with the method used for staining, provide relevant reagent descriptions as listed in MIFlowCyt guidelines (Section 2.4 Fluorescence Reagent(s) Descriptions). | For assessment of EV distribution in Optiprep density gradients, EVs were fluorescently labeled prior to density gradient centrifugation with 30 µM CFSE (Invitrogen, Carlsbad, CA) for 60 min at RT, after which the reaction was quenched by adding 2:1 volume:volume medium containing 10% EV-depleted FBS to the sample. |
| 2.2 Sample washing details | State any steps relating to the washing of samples. | No washing was performed. Unbound dye was inherently removed in CFSE experiments due to floatation in Optiprep density gradients. |
| 2.3 Sample dilution details | All methods and steps relating to sample dilution. | For assessment of EV distribution in Optiprep density gradients, individual gradient fractions 4-11 were diluted 1:20 in PBS immediately prior to flow cytometric measurements. For assessment of EV concentration in purified EV-preparations, pellets from pooled gradient fractions 6-9 were resuspended in 100 µL PBS and diluted a further 1:400 in PBS immediately prior to flow cytometric measurements. Samples were vortexed thoroughly after dilution and immediately prior to flow cytometric measurements. |
| 3.1 Buffer alone controls. | State whether a buffer-only control was analyzed at the same settings and during the same experiment as the samples of interest. If utilized it is recommended that all samples be recorded for a consistent set period of time e.g. 5 minutes, rather than stopping analysis at a set recorded event count e.g. 100,000 events. This allows comparisons of total particle counts between controls and samples. | A buffer-only control (PBS) was recorded using the same acquisition settings as all other samples during every measurement session. EV-concentrations in sample conditions were corrected for the background detected in the EV-gate for the buffer-only control in the same session. Mean EV-counts/uL for the buffer-only control was 307.8. |
| 3.2 Buffer with reagent controls. | State whether a buffer with reagent control was analyzed at the same settings, same concentrations, and during the same experiment as the samples of interest. If used state what the results were. | Reagent-only controls were not applied in this study. Throughout our previous work we have extensively tested the protocols used here, in terms of EV-depletion from FBS, CFSE staining and density gradient purification, which showed the ammount of carry-over of FBS EVs, lipoprotein particles and CFSE aggregates is negligible. |

| | | |
|---|---|---|
| 3.3 Unstained controls. | State whether unstained control samples were analyzed at the same settings and during the same experiment as stained samples. If used, state what the results were, preferably in standard units. | Unstained and stained samples were not measured side-by-side in this study. |
| 3.4 Isotype controls. | The use of isotype controls is applicable to immunofluorescence labelling only. State whether isotype controls were analyzed at the same settings and during the same experiment as stained samples. If utilized, state which antibody they are matched to, the concentration used, and what the results were (Section 4.2, 4.3, 4.4). Due to conjugation differences between manufacturers if should be stated if the isotype controls are from the same manufacturer as the matched antibodies. | Immunofluorescent labeling was not performed in the flow cytometry experiments described in this study. |
| 3.5 Single-stained controls. | State whether single-stained controls were included. If used state whether the single-stained controls were recorded using the same settings, dilutions, and during the same experiment as stained samples and state what the results were, preferably in standard units (Section 4.2, 4.3, 4.4). | Immunofluorescent labeling was not performed in the flow cytometry experiments described in this study. |
| 3.6 Procedural controls. | State whether procedural controls were included. If used, state the procedure and if the procedural controls were acquired at the same settings and during the same experiment as stained samples. | Procedural controls were not included in this study. |
| 3.7 Serial dilutions. | State whether serial dilutions were performed on samples and note the dilution range and manner of testing. The fluorescence and/or scatter signal intensity would ideally be reported in standard units (see Section 4.3, 4.4) but arbitrary units can also be used. This data is best reported by plotting the recorded number events/concentration over a set period of time at different sample dilution. The median fluorescence intensity at each of the dilutions should also ideally be plotted on the same or a separate plot. | Serial dilutions were tested, which showed a linear correlation between dilution factor and event count, indicating swarm detection was absent. |
| 3.8. Detergent treated EV-samples | State whether samples were detergent treated to assess lability. If utilized, state what detergent was used, the end concentration of the detergent, and what the results were of the lysis. | EV samples were not detergent-treated for flow cytometry experiments. |
| 4.1 Trigger Channel(s) and Threshold(s). | The trigger channel(s) and threshold(s) used for event detection. Preferably, the fluorescence calibration (Section 4.3) and/or scatter calibration (Section 4.4) should be used in order to report the trigger channel(s) and threshold(s) in standardized units. | For assessment of EV distribution in Optiprep density gradients, detection was triggered on the B2 fluorescence detector . For assessment of EV concentration in purified EV-preparations, detection was triggered on a SSC threshold. Absolute thresholds can be found in the Materials & Methods of the associated publication. Fluorescence/scatter calibration was not performed because the goal was to assess relative differences in EV-concentrations between samples to facilitate normalization in follow-up experiments, rather than absolute quantification. |
| 4.2 Flow Rate / Volumetric quantification. | State if the flow rate was quantified/validated and if so, report the result and how they were obtained. | Samples were measured at the low flow rate setting, corresponding to 10-15 uL/min as measured by the instrument flow rate sensors. The flow rate was not externally validated in this study. The machine was set to measure events from exactly 20 µL, for assessment of EV distribution in Optiprep density gradients, or 5 µL, for assessment of EV concentration in purified EV-preparations. |

| | | |
|---|---|---|
| 4.3 Fluorescence Calibration. | State whether fluorescence calibration was implemented, and if so, report the materials and methods used, catalogue numbers, lot numbers, and supplied reference units for the standards. Fluorescence parameters may be reported in standardized units of MESF, ERF, or ABC beads. The type of regression used, and the resulting scatter plot of arbitrary data vs standard data for the reference particles should be supplied. | Fluorescence calibration was not performed because the goal was to assess relative differences in EV-concentrations between samples to facilitate normalization in follow-up experiments, rather than absolute quantification. |
| 4.4 Light Scatter Calibration. | State whether and how light scatter calibration was implemented. Light scatter parameters may be reported in standardized units of nm2, along with information required to reproduce the model. | Scatter calibration was not performed because the goal was to assess relative differences in EV-concentrations between samples to facilitate normalization in follow-up experiments, rather than absolute quantification. |
| 5.1 EV diameter/surface area/volume approximation. | State whether and how EV diameter, surface area, and/or volume has been calculated using FC measurements. | EV-diameter, -surface area and -volume were not approximated in this study. |
| 5.2 EV refractive index approximation. | State whether the EV refractive index has been approximated and how this was done. | EV-refractive index was not approximated by flow cytometry in this study. |
| 5.3 EV epitope number approximation. | State whether EV epitope number has been approximated, and if so, how it was approximated. | EV-epitope number was not approximated in this study. |
| 6.1 Completion of MIFlowCyt checklist. | Complete MIFlowCyt checklist criteria 1 to 4 using the MIFlowCyt guidelines. Template found at www.evflowcytometry.org. | Completed MIFlowCyt checklist is provided with the publication. |
| 6.2 Calibrated channel detection range | If fluorescence or scatter calibration has been carried out, authors should state whether the upper and lower limits of a calibrated detection channel were calculated in standardized units. This can be done by converting the arbitrary unit scale to a calibrated scaled, as discussed in Section 4.3 and 4.4, and providing the highest unit on this scale and the lowest detectable unit above the unstained population. The lowest unit at which a population is deemed 'positive' can be determined a variety of ways, including reporting the 99th percentile measurement unit of the unstained population for fluorescence. The chosen method for determining at what unit an event was deemed positive should be clearly outlined. | Fluorescence and scatter calibration were not implemented in this study. |
| 6.3 EV number/concentration. | State whether EV number/concentration has been reported. If calculated, it is preferable to report EV number/concentration in a standardized manner, stating the number/concentration between a set detection range. | EV-concentrations were determined and are shown in the publication's figures, either as events/uL (corrected for dilution factors) or fold change to control conditions, as described in the paper. |
| 6.4 EV brightness. | When applicable, state the method by which the brightness of EVs is reported in standardized units of scatter and/or fluorescence. | EV-brightness is not reported in this study. |
| 7.1. Sharing of data to a public repository. | Provide a link to the experimental data in a public data repository. | N.A. |
